# Dynamic adult tracheal plasticity drives stem cell adaptation to changes in intestinal homeostasis

**DOI:** 10.1101/2021.01.10.426079

**Authors:** Jessica Perochon, Yachuan Yu, Gabriel N. Aughey, Tony D. Southall, Julia B. Cordero

## Abstract

Coordination of stem cell function by local and niche-derived signals is essential to preserve adult tissue homeostasis and organismal health. The vasculature is a prominent component of multiple stem cell niches. However, its role in adult intestinal homeostasis remains largely understudied. Here, we uncover a previously unrecognised crosstalk between adult intestinal stem cells (ISCs) in *Drosophila* and the vasculature-like tracheal system, which is essential for intestinal regeneration. Following damage to the intestinal epithelium, gut-derived reactive oxygen species (ROS) activate tracheal HIF-1α and bidirectional FGF/FGFR signaling, leading to reversible remodelling of gut-associated terminal tracheal cells and ISC proliferation following damage. Unexpectedly, ROS-induced adult tracheal plasticity involves downregulation of the tracheal specification factor *trachealess* (*trh*) and upregulation of IGF2 mRNA-binding protein (IGF2BP2/Imp). Our results reveal a novel intestine/vasculature interorgan communication program, which is essential to adapt stem cells response to the proliferative demands of the intestinal epithelium.

## Main

Adult intestinal plasticity is largely owed to the action of stem cells, which must respond to constant signals from the intestinal epithelium and its microenvironment, to fulfil global tissue demands^1–3^. Surprisingly, little is known about the role of the vascular microenvironment in adult intestinal homeostasis.

The *Drosophila* tracheal system is an oxygen-delivering interconnected tubular network, functionally analogous to the mammalian vascular and respiratory systems^4^. Following specification from epidermal cells and the formation of a tracheal sac in the embryo, tracheal cells undergo extensive cell rearrangements and cell shape changes, leading to the formation of multicellular tubes that ramify into progressively thinner branches, culminating with a terminal tracheal cell^5^. *Drosophila* terminal tracheal cells (TTCs), analogous to mammalian vascular tip cells^6^, extend prominent cytoplasmic projections, which supply oxygen to their target tissues^4, 5, 7, 8^. While tracheal development and post-embryonic plasticity have been significantly studied in *Drosophila*^5, 9, 10^, there is scarce knowledge on the role and regulation of the adult tracheal system.

The adult *Drosophila* midgut shares remarkable homology with the mammalian intestine^11^. Critically, the midgut epithelium is maintained by intestinal stem cells (ISCs), which self-renew and replenish the differentiated intestinal lineage —secretory enteroendocrine cells and absorptive enterocytes— through the production of undifferentiated enteroblasts^12, 13^. As its mammalian counterpart, the *Drosophila* gastrointestinal tract is densely tracheated^14^. Beyond the requirement for tracheal derived Dpp/BMP to restrain ISC proliferation^15^, there is no knowledge on the role of the tracheal system in adult midgut biology.

Here, we combine genetics and image analysis with *in vivo* functional and molecular studies to characterise a novel inter-organ communication program between the adult *Drosophila* midgut and its closely associated tracheal tissue, which is essential to shape stem cell and tracheal plasticity during intestinal regeneration.

## Results

### Intestinal damage leads to dynamic and reversible remodelling of gut associated adult TTCs in *Drosophila*

The adult *Drosophila melanogaster* gut is extensively covered by TTCs, labelled with a *GAL4* reporter driven under the control of the *Drosophila* homologue of Serum Response Factor (dSRF)^16–18^ (*dSRF>GFP*) (Fig. 1a and Extended data Fig. 1a). Transmission electron microscopy of adult posterior midguts denoted intimate contact between TTCs, enterocytes (ECs) (Fig. 1b) and ISCs (Fig. 1c). Oxygen and nutrient availability are two recognised determinants of TTC plasticity^10, 19^. We noticed that, damage to the adult *Drosophila* midgut epithelium caused by feeding animals with the pathogenic bacteria *Pseudomonas entomophila* (*Pe*)^20–23^, the DNA-damaging agent Bleomycin^24–26^, or the epithelial basement membrane disruptor Dextran Sulfate Sodium (DSS)^25, 27^, led to a significant increase in TTC coverage within the posterior midgut (Fig. 1d, e and Extended data Fig. 1b). Consideration of gut resizing upon *Pe* damage did not impact the overall increase in tracheal coverage (Extended data Fig. 1c-e). GFP labelled single terminal tracheal cell clones, unambiguously confirmed the increase in total number of cellular branches derived from individual TTCs in damaged (*Pe*) versus control (Sucrose) midguts (Fig. 1f, g). Interestingly, observation of single TTC clones within *Pe* treated midguts revealed direct correlation between the number of individual TTC branches and nearby PH3^+^ ISCs (Fig. 1h). Further quantification of tracheal phenotypes showed increase in primary, secondary and tertiary tracheal branches and total length of individual TTC extensions in damaged (*Pe*) versus control (Sucrose) midguts (Fig. 1i-k). Quantification of TTC numbers or assessment of potential co-localization between TTC bodies and the cell proliferation marker PH3 revealed no evidence of TTC proliferation following midgut damage (Extended data Fig. 1f, g). Collectively, these data suggest extensive cellular remodelling of TTCs in response to epithelial intestinal injury. A time course assessment of posterior midguts over a 16 hrs period of *Pe* infection (Damage phase) followed by 32 hrs on normal diet (Recovery phase) revealed direct correlation between tracheal coverage and ISC proliferation (Fig. 1l, m and Extended data Fig. 1h, i). These results strongly suggest that adult gut-associated-tracheal remodelling is a highly dynamic and reversible process, which accompanies changes in intestinal homeostasis.

**Fig. 1.**
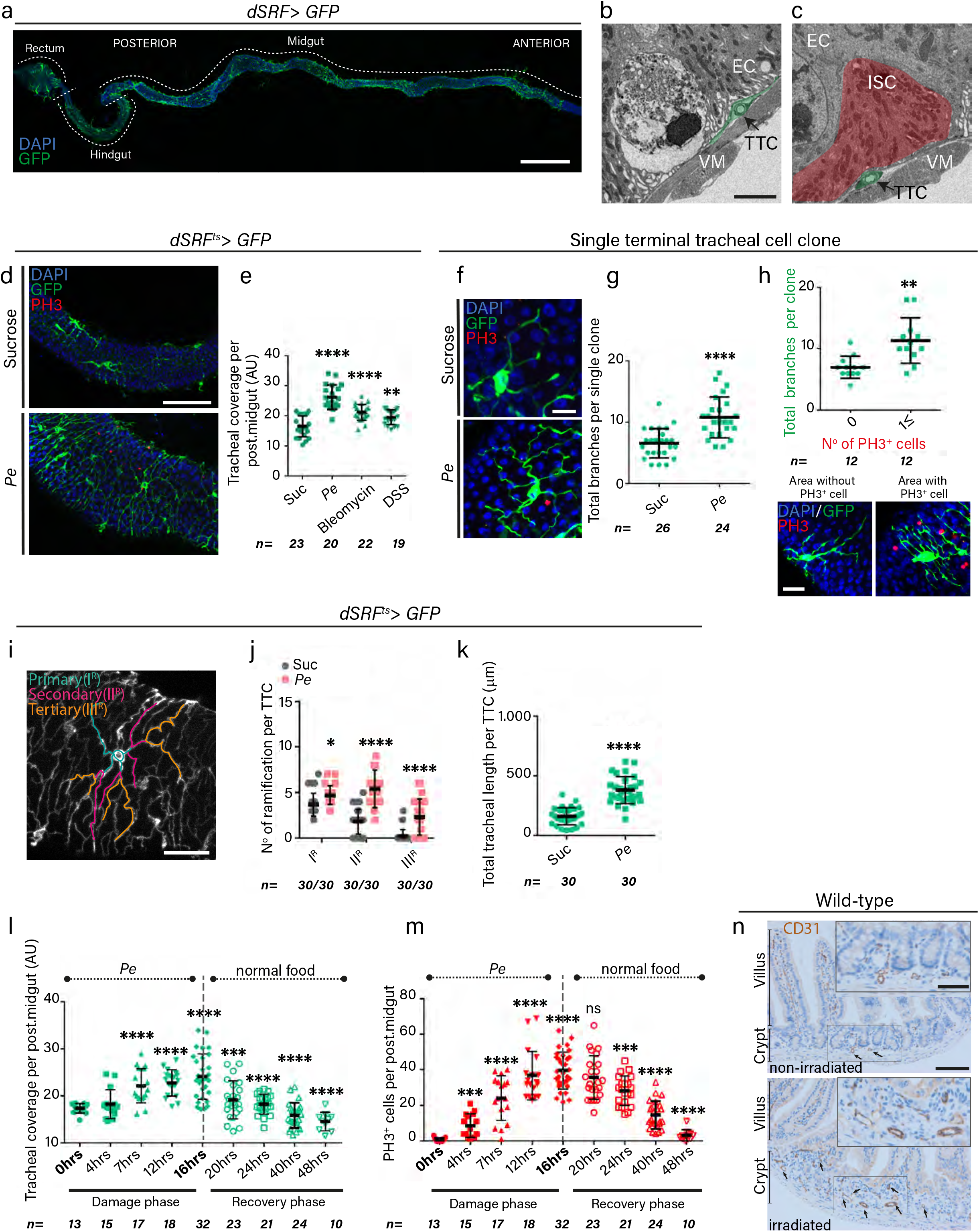
Adult tracheal plasticity following intestinal injury. **a**, Confocal image of adult gut-associated terminal tracheal cells (TTCs) (green) in *Drosophila*. Scale bar: 500μm. **b**, **c**, Transmission electron microscopy of adult posterior midguts highlighting TTCs (green pseudo-coloured), enterocyte (EC) and intestinal stem cell (ISC) (red pseudo-coloured); VM: visceral muscle, Scale bars: 5μm. **d**, Confocal images of TTCs (green) and proliferating ISCs (PH3; red) in control (Sucrose) or damaged (*Pe*) midguts. Scale bars: 100μm. **e,** Quantification of tracheal coverage in midguts as in (d) or upon Bleomycin or DSS treatment. **f**, Representative confocal images of single TTC clones (green) from control (Suc) or *Pe* damaged posterior midguts and proliferating ISCs (PH3; red). Scale bar: 25μm. **g**, Quantification of the total number of branches per TTC clone in midguts as in (f). **h, (top),** Quantification of total number of branches per TTC clone and nearby PH3^+^ ISCs upon *Pe* infection. **(bottom),** Representative confocal images of TTC clones (green) and proliferating ISCs (PH3; red) used for quantifications in top panel. Scale bars: 25μm. **f-h** <u>Statistics</u>: Student’s t test; n= number of TTC clones from 7-9 posterior midguts. **i,** Confocal image of a TTC with pseudo-coloured primary (I^R^), secondary (II^R^) and tertiary (III^R^) branches. Scale bar: 50μm. **j**, Quantification of number of branches per TTC from control or *Pe* infected posterior midguts. <u>Statistics</u>: Two-way ANOVA followed by Sidak’s multiple comparisons test; n= number of TTCs from 6 midguts per condition. **k**, Quantification of total tracheal length per TTC. Statistics: Student’s t test; n= number of TTCs from 6 midguts per condition. **l**, **m**, Quantification of tracheal coverage and PH3^+^ cells in posterior midguts during 16 hrs of oral *Pe* infection followed by 32 hrs standard food incubation. <u>Statistics</u>: Student’s t test to compare each timepoint of the damage and recovery stage against the 0 hrs and 16 hrs time points, respectively. n= number posterior midguts. **n**, Immunohistochemical analysis of control (non-irradiated) and regenerating (irradiated) mouse small intestines stained with anti-CD31 to visualize endothelial cells. Scale bars: 100μm (main figure); 50μm (close up view). Error bars: ± SEM, *p < 0.05, **p < 0.01, ***p < 0.001 ****p< 0.0001.

Low doses of whole body γ-irradiation in mice induce intestinal epithelial cell death, followed by a strong peak of crypt cell proliferation between 72- and 96 hrs after irradiation^28^. Staining with anti-CD31, showed an increase in vascular endothelial cells in regenerating (irradiated) versus control (non-irradiated) intestinal crypts (Fig. 1n). These results indicate a conserved phenomenology of vasculature/tracheal response to damage in the adult intestinal epithelium.

### TTC remodelling is necessary for ISC proliferation following damage of the adult *Drosophila* midgut

An essential step in the intestinal regenerative response to damage, involves a robust increase in ISC proliferation^1, 20 22, 25, 29^ (Fig. 1d, h). To address the functional role of the tracheal system in adult intestinal regeneration, we severely reduced trachea by overexpressing the pro-apoptotic gene *bax* (*UAS-bax*) in adult TTCs using temperature sensitive *dSRF-Gal4* (*dSRF^ts^>bax*) (Fig. 2a, b). This caused a significant impairment of the regenerative response of the intestine to multiple damages, as evidenced by an approximate 50% decrease in damage induced ISC proliferation (Fig. 2c). TTCs are best known for their role facilitating gas exchange with their target tissues^4^. Thus, poor intestinal regeneration following TTC reduction might reflect the need of oxygen in this process. Exposing animals to high and prolonged hypoxic environmental conditions, induced activity of a reporter of *Drosophila* Hypoxia-inducible factor-1α (HIF-1α/Sima)^30^ and TTC remodelling in the adult midgut (Extended data Fig. 2a-c). However, while hypoxia clearly impaired damage induced ISC proliferation in the adult midgut (Fig. 2d), it did so to a lower extent than that observed upon TTC loss (*dSRF^ts^>bax*) (Fig. 2c). This difference could be due to a compensatory effect of increased trachea upon hypoxia (Extended data Fig. 2a, c) or compromised ISC survival in *dSRF^ts^>bax* midguts. We assessed apoptosis and ISC numbers in hypoxic and *dSRF^ts^>bax* midguts through anti-caspase (Dcp-1) staining and the use of an ISC reporter (*Delta-LacZ*). Midguts overexpressing *bax* in adult ECs (*NP1^ts^>bax*), served as a ‘cell-death’ positive control (Extended data Fig. 2d, f). While we saw no evidence of cell death in hypoxic midguts, *dSRF^ts^>bax* midguts showed significant apoptosis (Extended data Fig. 2e, g), which was restricted to ECs, distinguished by their large nuclei (Extended data Fig. 2e, lower panel, magnified view). Consistently, this cell death phenotype did not translate into defective ISC numbers (Extended data Fig. 2h, i). Therefore, impaired midgut regeneration following the hypoxia or TTC ablation regime used in our study is unlikely to be secondary to ISC loss. Alternatively, differences in the regenerative response of hypoxic versus *dSRF^ts^>bax* midguts could be explained by the contribution of angiocrine factors to ISC proliferation, in addition to oxygen availability. Dpp/BMP ligand has been identified as an angiocrine factor in the adult midgut^15^. However, its action inhibits rather than induces ISC proliferation^15^. Therefore, a potential role of Dpp cannot directly explain our results (see also Discussion).

**Fig. 2.**
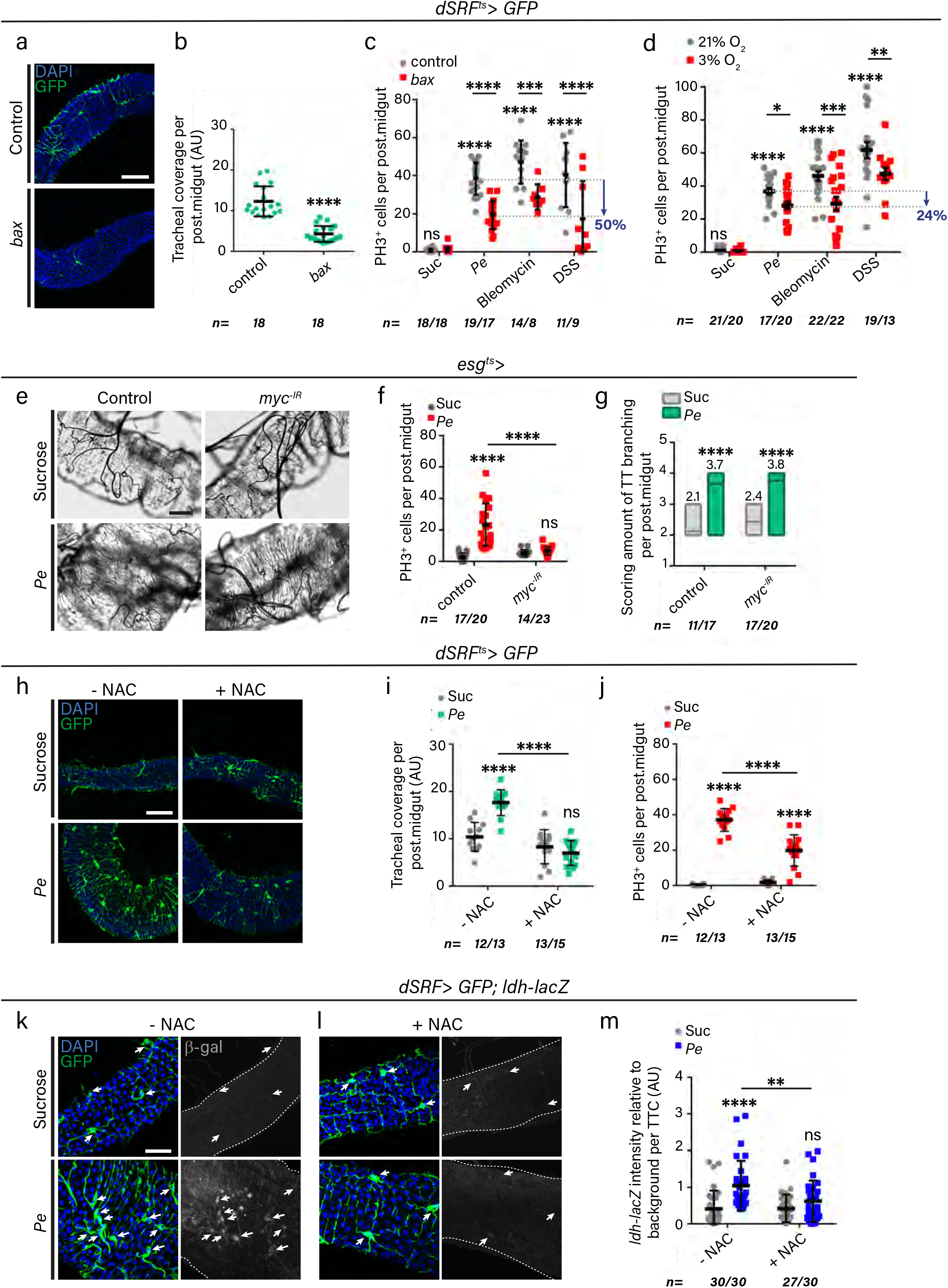
ROS induced tracheal remodelling is required for intestinal regeneration. **a,** Confocal images of adult posterior midguts from control animals or upon adult-specific *bax-* driven TTC loss (green). Scale bar: 100μm. **b**, Quantification of tracheal coverage in posterior midguts as in (a). <u>Statistics</u>: Student’s t test; n= number of posterior midguts. **c-d**, Quantification of PH3^+^ ISCs in posterior midguts of the genotypes and conditions indicated. **e**, Representative brightfield images of adult posterior midguts from Sucrose fed and *Pe* infected animals without (control) or with adult-specific overexpression of *myc* RNAi in ISCs/EBs (*esg^ts^>*). Scale bar: 50μm. **f**, **g**, Quantification of PH3^+^ ISCs (f) and scored tracheal branching (g) in posterior midguts as in (e). **h**, Confocal images of control and *Pe* treated midguts in the presence or absence of the antioxidant NAC. Scale bar: 100μm. **i**, **j**, Quantification of tracheal coverage (i) and PH3^+^ ISCs (j) in midguts as in (h). **c-j,** <u>Statistics</u>: Two-way ANOVA followed by Sidak’s multiple comparisons test; n= number of posterior midguts. **k**,**l,** Confocal images of the HIF-1α/Sima activity reporter *ldh-lacZ* (grey) in control or *Pe* infected adult posterior midguts without or with NAC (k, l, respectively). Scale bar: 50μm. **m**, Quantification of the average *ldh-lacZ* staining intensity in TTC within a defined region of posterior midguts as in (k, l). <u>Statistics</u>: Two-way ANOVA followed by Sidak’s multiple comparisons test; n= number of TTC quantified from 9 to 10 posterior midguts per condition. Error bars: ±SEM; *p < 0.05, **p < 0.01, ***p < 0.001 ****p< 0.0001.

Given that tracheal remodelling and ISC proliferation show almost the exact same dynamics (Fig. 1l, m and Extended data Fig. 1 h, i), it is unclear whether these events are part of a feedforward mechanism or if one precedes the other. To address this, we assessed tracheal remodelling following damage while blocking ISC proliferation by overexpressing *UAS-myc RNAi* (*myc^-IR^*)^31^ using the stem/progenitor driver *escargot-GAL4* (*esg^ts^>myc^-IR^*) (Fig. 2e-g). As we needed to use the *GAL4/UAS* system to genetically manipulate gut cells, we established a scoring method for assessing tracheal coverage through the use of light microscopy, which was validated against our confocal microscopy tracheal quantification approach (Extended Data Fig. 3a-c). Gut-associated trachea remodelled normally in *esg^ts^>myc^-IR^* midguts following *Pe* damage, in spite of the almost complete absence of ISC proliferation (Fig. 2e-g). Therefore, TTC remodelling precedes midgut ISCs proliferation following damage. We hypothesised that signals activated by damage upstream of ISC proliferation might induce gut-tracheal remodelling.

### Gut-derived ROS initiates tracheal remodelling through activation of HIF-1α/FGFR signaling in TTCs

Pathogen-induced intestinal damage triggers a strong oxidative burst and the production of reactive oxygen species (ROS) from the intestinal epithelium^32, 33^. We therefore tested whether ROS could trigger tracheal remodelling in the regenerating intestine. Feeding animals with the antioxidant N-acetyl cysteine (NAC) or genetically blocking ROS production in ECs by overexpressing the enzyme *catalase*^32^ (*NP1^ts^>catalase*) significantly impaired damage-induced tracheal remodelling (Fig. 2h, i and Extended data Fig. 3d, e) in addition to the expected reduction in regenerative ISC proliferation (Fig. 2j and Extended data Fig. 3f)^32^. Conversely, driving adult intestinal epithelial cell death through *bax* overexpression in ECs (*NP1^ts^>bax*) was sufficient to induce TTC remodelling and ISC proliferation (Extended data Fig. 3g-i). Therefore, intestinal epithelial damage and ROS induce remodelling of gut associated trachea, which is in turn necessary to drive ISC proliferation during intestinal regeneration.

Exogenous H_2_O_2_ can stabilize HIF-1α —a key conserved driver of hypoxia-induced tracheal/vascular remodelling^9, 34, 35^— in normoxia^36^. The Sima/HIF-1α activity reporter *ldh-lacZ* was upregulated in gut-associated TTCs following midgut damage and in an ROS dependent manner (Fig. 2k-m). Furthermore, midguts from *sima^-/-^* whole mutant animals or upon adult specific *sima* knockdown within TTCs (*dSRF^ts^>sima^-IR^*) showed significantly impaired tracheal remodelling and ISC proliferation following damage (Fig. 3a-f).

**Fig. 3.**
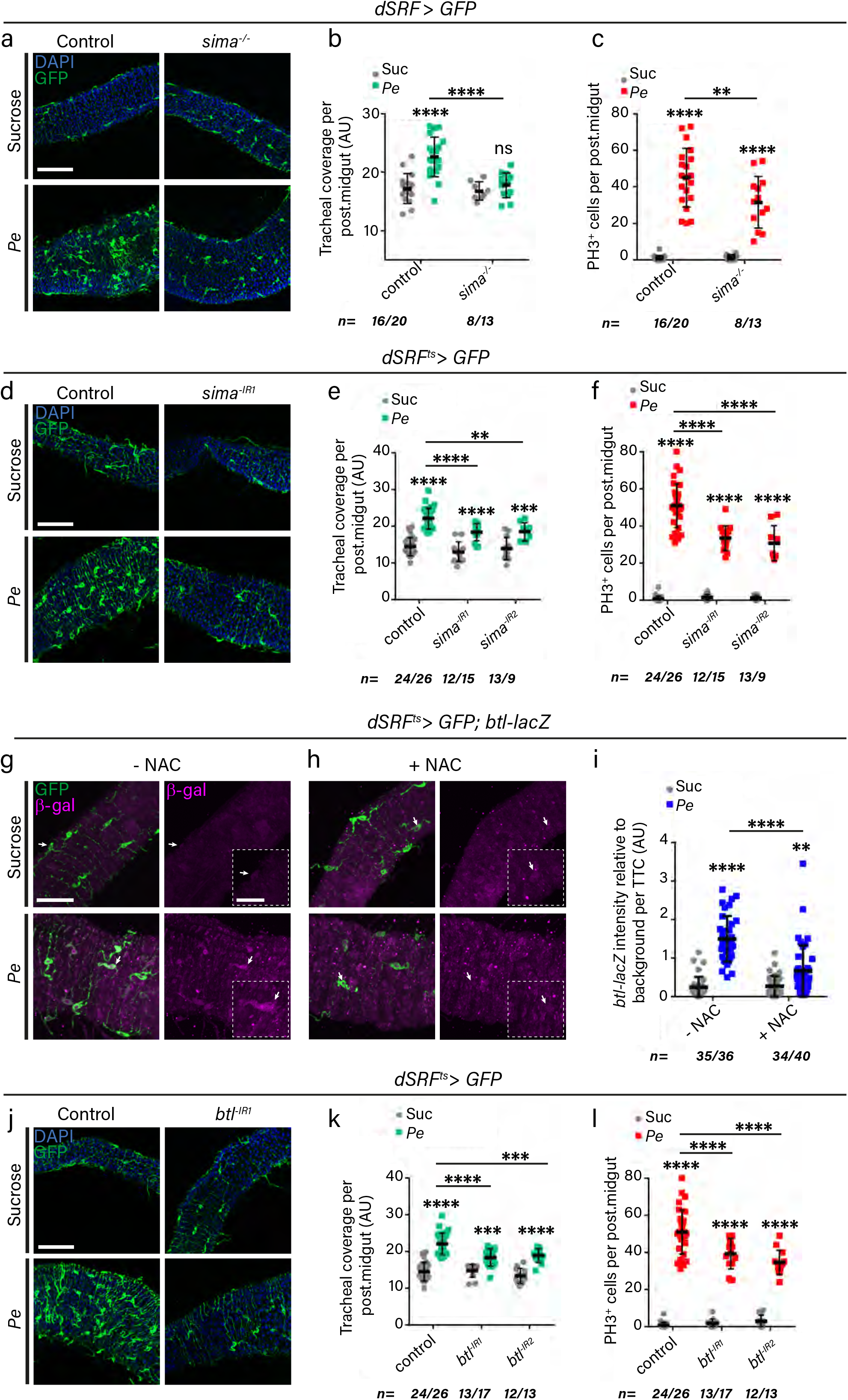
Activation of HIF-1α/FGFR signaling in TTCs is required for tracheal remodelling and intestinal regeneration following damage. **a,** Confocal images of TTCs (green) in control (Sucrose) or *Pe* infected adult posterior midguts from wild type or *Sima^-/-^* whole mutant animals. Scale bar: 100μm. **b**, **c**, Quantification of tracheal coverage (b) and PH3^+^ ISCs (c) in posterior midguts as in (a). **d**, Confocal images of control (Sucrose) or *Pe* infected adult posterior midguts from wild type animals or animals subject to adult-specific *sima* RNAi overexpression (*sima^-IR^*) within TTC (*dSRF^ts^>GFP*). Scale bar: 100μm. **e**, **f**, Quantification of tracheal coverage (e) and PH3^+^ ISCs in posterior midguts as in (d). **a-f,** <u>Statistics</u>: Two-way ANOVA followed by Sidak’s multiple comparisons test; n= number of posterior midguts. **g**, **h**, Representative confocal images of *btl* reporter expression (*btl-lacZ*; magenta) in control or *Pe* infected adult posterior midguts without or with NAC (g, h, respectively). Dotted boxes represent a higher magnification of the area pointed with an arrow. Scale bars: 50μm (main figure); 20μm (close up view). **i**, Quantification of *btl-lacZ* staining intensity in TTC relative to background within a defined region of posterior midguts as in (g, h). <u>Statistics</u>: Two-way ANOVA followed by Sidak’s multiple comparisons test; n= number of TTC from 9 to 10 posterior midguts/condition. **j**, Confocal images of control and damaged adult posterior midguts from wild type animals or following *RNAi*-driven adult-specific *btl* knockdown (*btl^-IR^*) within TTCs. Scale bar: 100μm. **k**, **l**, Quantification of tracheal coverage (k) and PH3^+^ ISCs in posterior midguts as in (j). <u>Statistics</u>: Two-way ANOVA followed by Sidak’s multiple comparisons test; n= number of posterior midguts. Error bars: ± SEM; *p < 0.05, **p < 0.01, ***p < 0.001 ****p< 0.0001.

The *Drosophila* fibroblast growth factor receptor (FGFR), Breathless (Btl), is a well-known transcriptional target of HIF-1α during tracheal development and oxygen-driven tracheal remodelling^19, 37^. Consistently, a reporter of *breathless (btl)* expression (*btl-lacZ)* showed gene upregulation in TTCs following intestinal damage, which was abrogated by NAC (Fig. 3g-i). TTC knockdown of *btl* (*dSRF^ts^>btl^-IR^*) inhibited tracheal remodelling and ISC proliferation following damage (Fig. 3j-l), without evidence of cell death or ISC loss (Extended data Fig. 2e, g-i). Therefore, ROS-dependent activation of HIF-1α/FGFR signaling within TTCs following gut damage induces tracheal remodelling and regenerative ISC proliferation in the adult midgut. Consistently, expression of the HIF-1α/FGFR target gene *blistered* (*bs*)/*dSRF*^19, 38^ was upregulated in damaged and hypoxic midguts (Extended data Fig. 4a-e) and knocking down *bs* in adult TTCs (*dSRF^ts^>bs^-IR^*) impaired tracheal remodelling and ISC proliferation following damage (Extended data Fig. 4f-h).

### ROS-dependent bidirectional FGF/FGFR signaling drives stem cell proliferation and TTC remodelling during intestinal regeneration

During development or hypoxia, the *Drosophila FGF*-like ligand Branchless (Bnl) is upregulated in target tissues and signals paracrinally to its receptor FGFR/Breathless (Btl) in the trachea to induce their remodelling^16, 19, 39^. Consistently, we observed upregulation of a *bnl* reporter (*bnl-lacZ*) in ISCs/EBs and ECs following intestinal damage, which was impaired by NAC (Fig. 4a-c). These results suggest that ROS induces Bnl activation within the intestinal epithelium following damage. Unexpectedly, using the same reporter, we observed that *bnl* was also upregulated in TTCs following intestinal damage, in an ROS dependent manner (Fig. 4d-e). Expression of *bnl* in TTCs was confirmed by the use of an independent reporter (Extended data Fig. 5a, b). Overexpressing *bnl* in adult TTCs (*dSRF^ts^>bnl*) was sufficient to induce ISC proliferation without TTC remodelling (Extended data Fig. 5c, d).

**Fig. 4.**
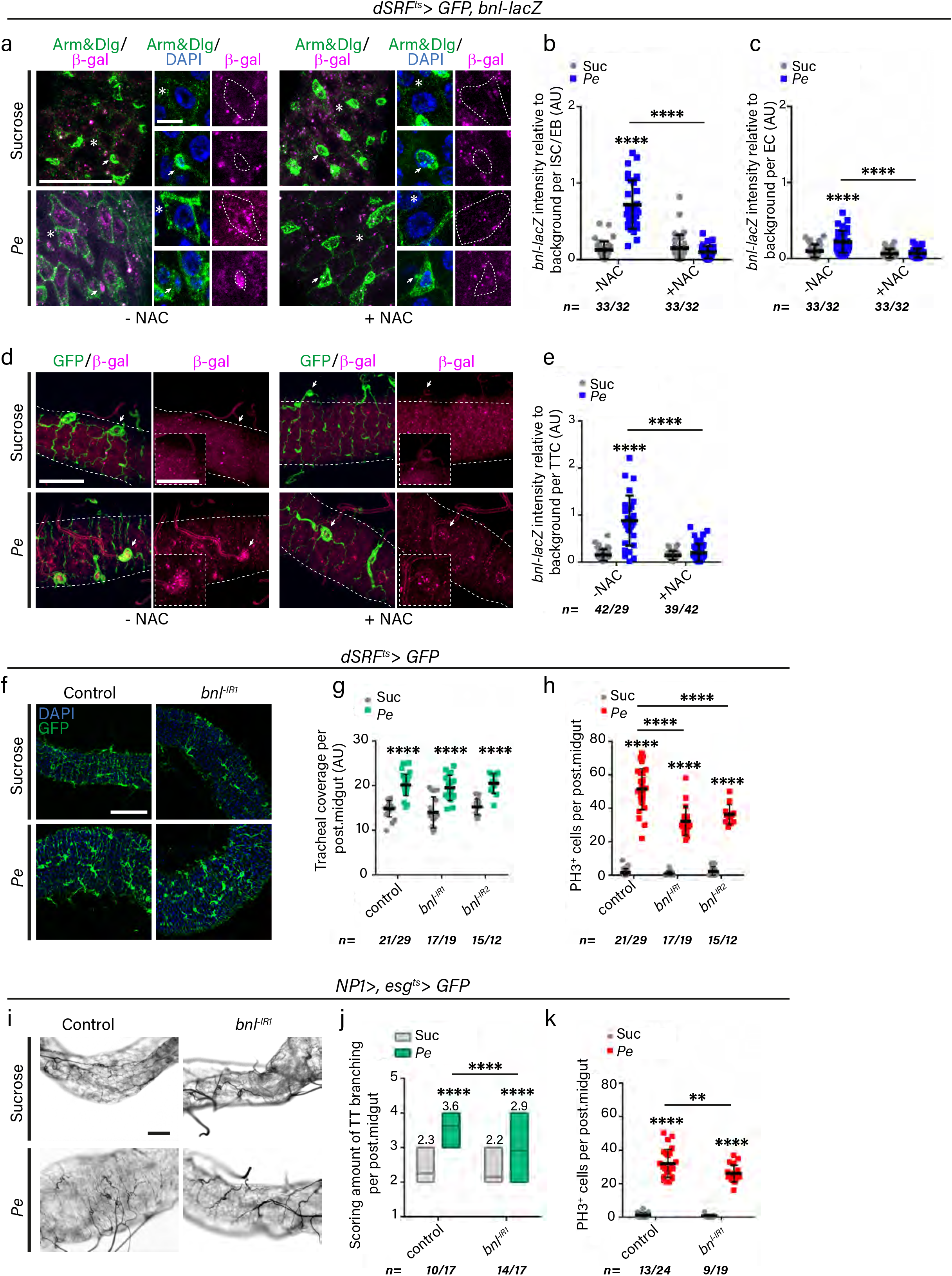
Bidirectional FGF/FGFR signaling between the midgut and TTCs drives tracheal remodelling and ISC proliferation during intestinal regeneration. **a,** Representative confocal images of *FGF/bnl* reporter expression (*bnl-lacZ*; magenta) in control and *Pe* treated midguts in the presence or absence of NAC. Arrows point to reporter signal within ISCs/EBs (small nuclei and stained with anti-Armadillo (Arm); pseudo-coloured in green). Asterisks point to reporter signal within ECs (large nuclei and stained with anti-Discs large (Dlg); pseudocoloured in green). Dotted areas outline ISCs/EBs or ECs pointed by arrows or asterisks, respectively. We noted consistent, unexplained downregulation of Dlg relative to Arm in Sucrose and NAC treated midguts. Scale bars: 40μm (main figure); 9μm (close up view). **b**, **c**, Quantification of *bnl-lacZ* staining within ISCs/EBs (b) and ECs (c) in posterior midguts as in (a). **d**, Representative confocal images of *FGF/bnl* reporter expression (*bnl-lacZ*; magenta) and TTCs (*dSRF>GFP*; green) in control (Sucrose) and regenerating (*Pe*) adult posterior midguts in the presence or absence of NAC. Arrows indicate reporter signal within TTCs. Dotted boxes show a magnified view of TTCs pointed by arrows. Scale bars: 40μm (main figure); 20μm (close up view). **e**, Quantification of *bnl-lacZ* staining within TTCs in posterior midguts as in (d). **a-e,** <u>Statistics</u>: Two-way ANOVA followed by Sidak’s multiple comparisons test; n= number of TTCs, ISC or ECs from 10-14 posterior midguts per condition. **f**, Confocal images of control and *Pe* treated adult posterior midguts from wild type animals or upon RNAi-driven adult-specific *bnl* knockdown (*bnl^-IR^*) within TTCs. Scale bar: 50μm. **g**, **h**, Quantification of tracheal coverage (g) and PH3^+^ ISCs (h) in midguts as in (f). **i**, Representative brightfield images of control (Sucrose) or regenerating (*Pe*) adult posterior midguts from wild type animals or upon RNAi-driven adult *bnl* knockdown (*bnl^-IR^*) within ISCs/EBs and ECs (*NP1>, esg^ts^>GFP*). Scale bar: 50μm. **j**, **k**, Quantification of scored tracheal branching (j) and PH3^+^ ISCs (k) in posterior midguts as in (i). **f-k,** <u>Statistics</u>: Two-way ANOVA followed by Sidak’s multiple comparisons test; n= number of posterior midguts. Error bars: ± SEM; *p < 0.05, **p < 0.01, ***p < 0.001 ****p< 0.0001.

We next assess the functional role of individual sources of FGF/Bnl in our system. Consistent with our reporter expression data (Fig. 4a-e), knocking down *bnl* from either TTCs (*dSRF^ts^>bnl^-IR^*), ISCs/EBs (*esg^ts^>bnl^-IR^*) or ECs (*NP1^ts^>bnl^-IR^*) restrained ISC proliferation following midgut damage but did not impair TTC remodelling (Fig. 4f-h and Extended data Fig. 5e-j). This is in line with the high sensitivity of the regenerative intestine to discrete fluctuations in individual signaling activity^31, 40^. Hence, small variations in Bnl levels, which are insufficient to affect tracheal remodelling are enough to impact ISC proliferation following damage. Instead, concomitant *bnl* knockdown from ECs and ISCs/EBs (*NP1>, esgts>bnl^-IR^*) significantly impaired TTC remodelling and subsequent ISC proliferation (Fig. 4i-k). Therefore, the combined action of gut-derived sources of Bnl, is necessary to induce tracheal remodelling following intestinal damage.

Given that multiple sources of Bnl—from the midgut and TTCs—can individually contribute to regenerative ISC proliferation independently of tracheal remodelling, we hypothesised this may be through a non-tracheal receptor. Consistently, knocking down *btl* from ISCs/EBs (*esg^ts^>btl-IR*) significantly impaired ISC proliferation upon damage without affecting TTC remodelling (Extended data Fig. 6a-c). In the context of tracheal development, Bnl/Btl signals through the MAPK/ERK pathway^5, 41^, which is a key driver of ISC proliferation in the adult *Drosophila* midgut^42–44^. Knocking down *btl* from ISCs/EBs (*esg^ts^>btl^-IR^*) impaired damage-induced MAPK/ERK activation in the midgut (Extended data Fig. 6d, e), suggesting that activation of Btl in the midgut regulates regenerative ISC proliferation through MAPK/ERK signaling.

### Identification of novel TTC intrinsic mechanisms triggered during intestinal regeneration

We next used Targeted DamID (TaDa) for TTC *in vivo* profiling of RNA Pol II chromatin binding^45^ in control (Sucrose) and *Pe* treated midguts (Fig. 5a and Extended data Fig. 7a). TaDa is particularly advantageous in our system due to inherent difficulties to efficiently separate tracheal tissue from the midgut. We identified 1747 and 1712 genes significantly bound by RNA Pol II in TTCs from control (Sucrose) and *Pe* infected midguts, respectively (Supplementary Table 1). Gene ontology (GO) analysis of areas with significant RNA Pol II binding in control midguts revealed enrichment in components of the tracheal system, and genes previously involved in epithelial tube morphogenesis and respiratory/tracheal system development (Fig. 5b) (Supplementary Table 2 and 3). This validated the sensitivity of TaDa to reliably detect tracheal specific genes from combined gut and tracheal tissue samples.

**Fig. 5.**
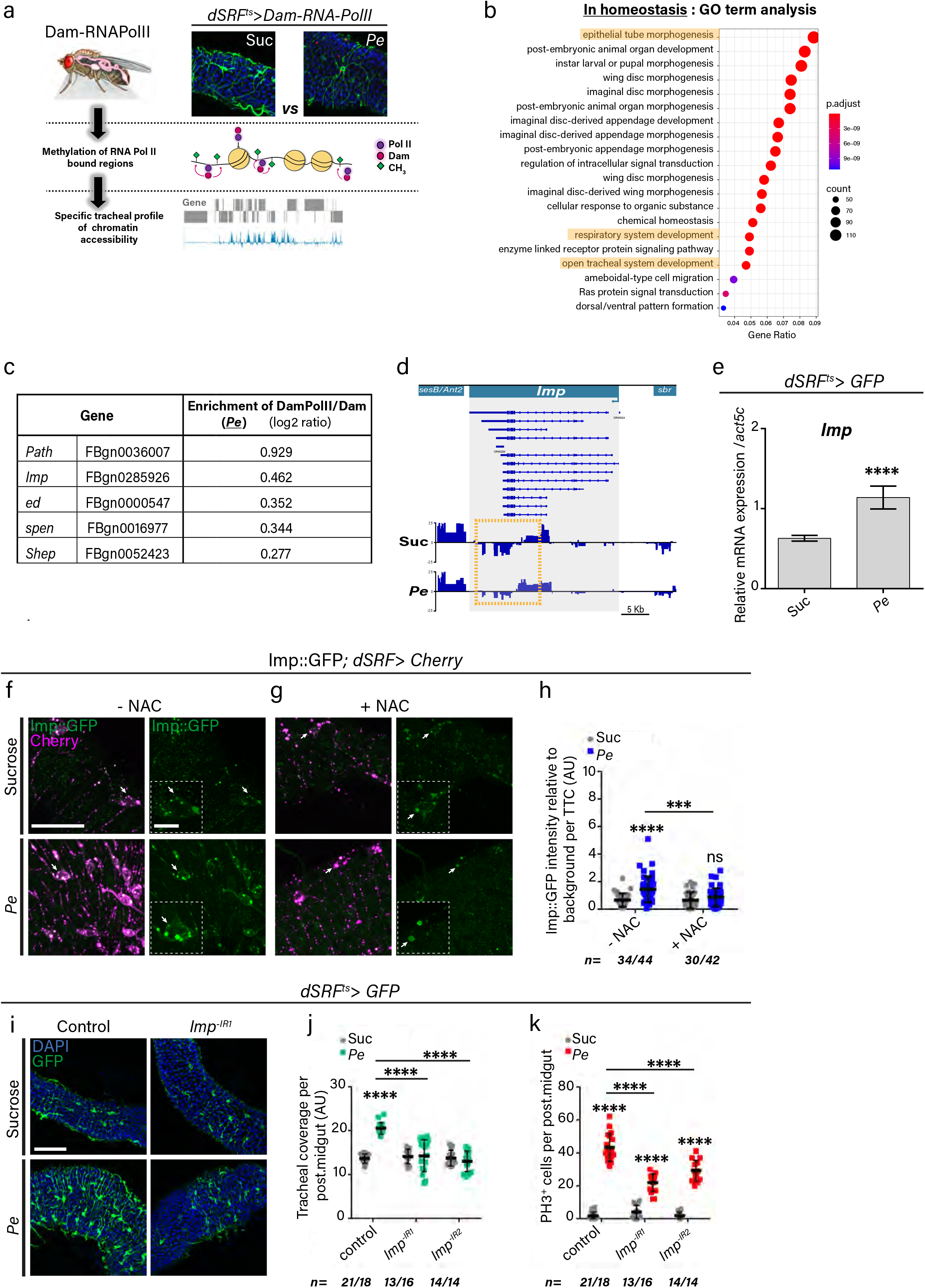
Imp is a novel regulator of adult tracheal remodelling and intestinal regeneration. **a,** Schematic representation of targeted DamID approach used to profile RNA pol II binding in adult TTCs. **b**, Gene Ontology term analysis corresponding to genes with significant RNA pol II binding in adult TTCs from control midguts (Sucrose). **c**, List of genes associated to neuronal processes showing significant RNA pol II binding in TTCs following *Pe* treatment. Data is presented as the Log2 ratio of Dam-RNA Pol II/Dam (normalization control) read counts. **d**, Profile of RNA Pol II binding to *Imp* in TTCs from control (Suc) and regenerating (*Pe*) midguts. Boxes highlight areas with most prominent differences in RNA Pol II binding. **e**, qRT-PCR for *Imp* mRNA expression from whole adult midguts and associated tracheal tissue, in control (Sucrose) or *Pe* fed wild type animals. <u>Statistics</u>: Data are represented as mean ± SEM of five biological replicates. Student’s t test. **f**, **g**, Representative confocal images of Imp protein reporter Imp::GFP (green) and TTCs (*dSRF>Cherry*; magenta) in control (Sucrose) and regenerating (*Pe*) posterior midguts in the absence (f) or presence (g) of NAC. Arrows indicate reporter signal within TTCs. Dotted boxes show magnified views of TTCs pointed by arrows. Scale bars: 50μm (main figure); 12μm (close up view). **h**, Quantification of Imp::GFP staining intensity relative to background within TTCs in posterior midguts as in (f) and (g). <u>Statistics</u>: Two-way ANOVA followed by Sidak’s multiple comparisons test; n= number of TTC from 11-14 posterior midguts per condition. **i**, Representative confocal images of control (Sucrose) or *Pe* treated adult posterior midguts from wild type animals or upon RNAi-driven adult-specific *Imp* knockdown (*Imp^-IR^*) within TTCs (green). Scale bar: 100μm. **j**, **k**, Quantification of tracheal coverage (j) and PH3^+^ ISCs (k) in posterior midguts as in (i). <u>Statistics</u>: Two-way ANOVA followed by Sidak’s multiple comparisons test; n= number of posterior midguts. Error bars: ± SEM; *p < 0.05, **p < 0.01, ***p < 0.001 ****p< 0.0001.

Consistent with our reporter expression and functional data (Fig. 4d-h), TaDa analysis identified *bnl/FGF* as a gene with significant RNA Pol II binding in *Pe* treated midguts only (Supplementary Table 1) (Extended Data Fig. 7b). Unexpectedly, we were unable to detect significant RNA Pol II binding to *btl* in adult TTCs of *Pe* treated midguts (Supplementary Table 1). This is counterintuitive given our gene expression and functional data on *btl* (Fig. 3g-l).

Discrepancies between RNA pol II occupancy and mRNA transcript status are possible and could be due to pausing of the polymerase^46^, post-transcriptional mRNA regulation^47^ or temporally dynamic RNA pol II binding (*e.g* during intestinal damage), which may not be captured by a single time point assessment.

### Imp/IGF2BP is a novel regulator of adult tracheal remodelling and intestinal regeneration

Interestingly, we noticed that, within the genes showing significant RNA pol II binding in TTCs of *Pe* treated midguts only (Supplementary Table 1), there were several genes associated with neuronal function (Fig. 5c). Amongst them, was the highly conserved mRNA-binding protein Imp/IGF2BP (Fig. 5d) (Supplementary Table 1), which regulates axonal remodelling in *Drosophila*^48, 49^. We confirmed upregulation of *Imp* transcription by qRT-PCR (Fig. 5e) and Imp increase in TTCs through the use of a protein trap (Imp::GFP) (Fig. 5f, h). NAC treatment showed that Imp upregulation in TTCs following intestinal damage depends on ROS production (Fig. 5g, h). Importantly, adult specific knock down of *Imp* from TTCs (*dSRF^ts^>Imp^-IR^*) significantly impaired tracheal remodelling and ISC proliferation following midgut epithelial damage (Fig. 5i-k). Next, we investigated two well-known post-transcriptional targets of Imp in our system: *chickadee/profilin*^48^ and *myc*^49^. Functional experiments on the role of *chickadee* led to inconclusive results (data not shown). However, Myc was upregulated in TTCs of *Pe* treated midguts and this was abrogated by *Imp* knockdown (*dSRF^ts^>imp^-IR^*) (Fig. 6a, b). Moreover, adult specific knockdown of *myc* within TTCs using RNAi^31^ (*dSRF^ts^>imp^-IR^*), impaired tracheal remodelling and ISC proliferation following gut damage (Fig. 6c-e). Altogether, these results establish Imp as a novel regulator of TTC remodelling and ISC proliferation during adult *Drosophila* midgut regeneration. This function of Imp is at least in part through tracheal intrinsic control of Myc.

**Fig. 6.**
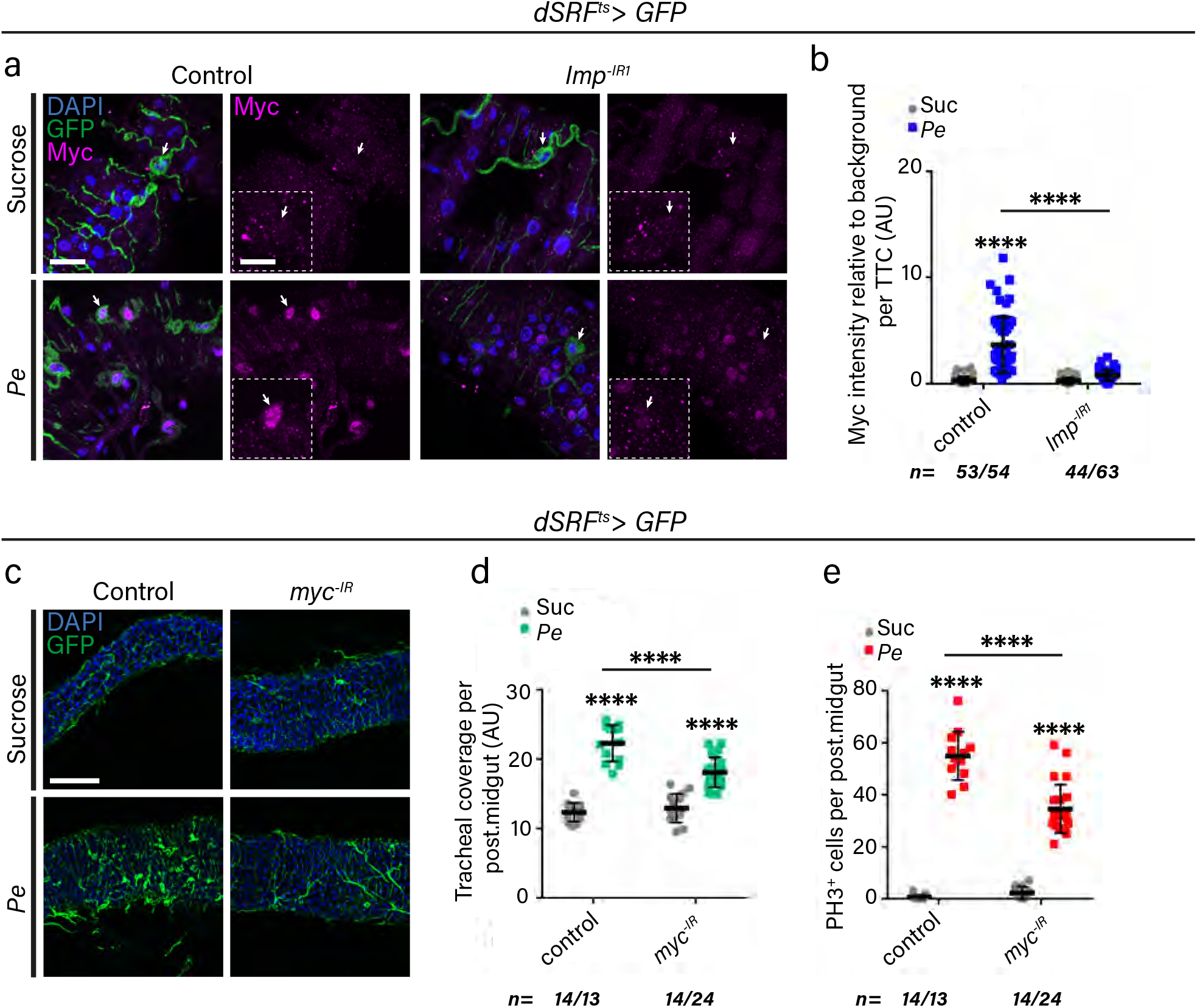
Imp dependent activation of Myc in TTCs is required for tracheal remodelling and intestinal regeneration. **a,** Representative confocal images of Myc staining (magenta) and TTCs (*dSRF>GFP*; green) in control (Sucrose) or regenerating (*Pe*) midguts from wild type animals or upon RNAi-driven adult-specific *Imp* knockdown (*Imp^-IR^*) within TTCs. Arrows indicate Myc signal in TTCs. Dotted boxes show magnified views of TTCs pointed by arrows. Scale bars: 20μm (main figure); 10μm (close up view). **b**, Quantification of Myc staining intensity relative to background in TTCs from posterior midguts as in (a). <u>Statistics</u>: Two-way ANOVA followed by Sidak’s multiple comparisons test; n= number of TTCs from 15-18 posterior midguts per condition. **c**, Representative confocal images of control (Suc) or *Pe* treated adult posterior midguts from wild type animals or upon RNAi-driven adult-specific *myc* knockdown (*myc^-IR^*) within TTCs (green). Scale bar: 100μm. **d**, **e**, Quantification of tracheal coverage (d) and PH3^+^ ISCs (e) in posterior midguts as in (c). <u>Statistics</u>: Two-way ANOVA followed by Sidak’s multiple comparisons test; n= number of posterior midguts quantified. Error bars: ± SEM; *p < 0.05, **p < 0.01, ***p < 0.001 ****p< 0.0001.

### *Trachealess* downregulation in TTCs is necessary for adult tracheal remodelling and damage induced ISC proliferation

While our TaDa analysis revealed significant binding of RNA Pol II to *trachealess* (*trh*) in TTCs of Sucrose treated midguts (Supplementary Table 1 and 3), this was not the case in the damaged tissues (Fig. 7a and Supplementary Table 1). This was surprising given that *trh* is known a master regulator of tracheal gene expression and it is present in all tracheal cells from the onset of embryonic development through adulthood^50–52^. Loss of *trh* during development impairs tracheal cell specification and tube morphogenesis^51, 52^. However, qRT-PCR confirmed downregulation of *trh* upon intestinal damage (Fig. 7b). Antibody staining (Fig. 7c, d) and a transgenic reporter (*trh-lacZ*) (Fig. 7e, f) further demonstrated protein and gene downregulation in TTCs upon gut damage, respectively. Remarkably, *trh-lacZ* signal was significantly restored upon *Pe* and NAC co-treatment or 32 hrs after removal of the damaging agent (Fig. 7e, f). This suggests that *trh* expression in adult TTCs is highly dynamic and its downregulation upon intestinal damage is dependent on ROS. Importantly, consistent with our gene and protein expression data, *trh* overexpression in adult TTCs (*dSRF^ts^>trh*) significantly impaired tracheal remodelling and ISC proliferation (Fig. 7g-i), while *trh* knockdown (*dSRF^ts^>trh^-IR^*) potentiates TTC remodelling and ISC proliferation following midgut damage (Fig. 7g-i). Altogether, these results suggest that ROS-induced *trh* downregulation in adult TTCs is necessary to allow gut associated TTC plasticity and robust regeneration of the intestine following damage.

**Fig. 7.**
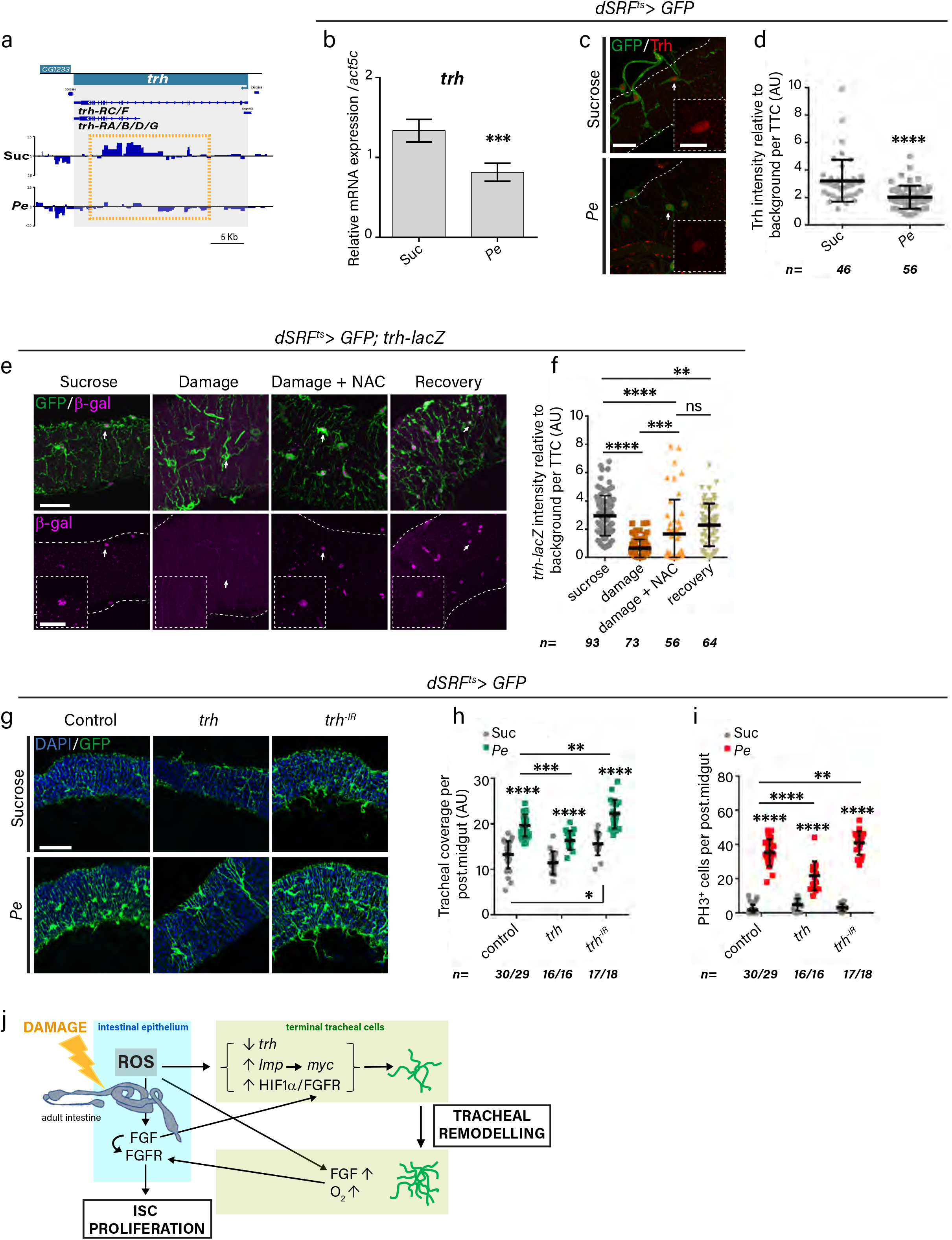
*Trachealess* downregulation in TTCs is necessary for adult tracheal remodelling and damage induced ISC proliferation. **a,** Profile of RNA Pol II binding to *trachealess* (*trh*) within TTCs of control (Suc) and regenerating (*Pe*) midguts. **b**, qRT-PCR for *trh* mRNA expression from whole adult midguts and associated tracheal tissue of control (Sucrose) or *Pe* fed animals. <u>Statistics</u>: Data are represented as mean ± SEM of five biological experimental replicates. Student’s t test. **c**, Representative confocal images of Trh staining (red) and TTCs (*dSRF>GFP*; green) in control (Sucrose) or regenerating (*Pe*) midguts. Arrows indicate Trh signal within TTCs. Dotted boxes show magnified views of TTCs pointed by arrows. Scale bars: 20μm (main figure); 10μm (close up view). **d**, Quantification of Trh staining intensity relative to background within TTCs in posterior midguts as in (c). <u>Statistics</u>: Student’s t test; n= number of TTC from 14 posterior midguts per condition. **e**, Representative confocal images of *trh* reporter expression (*trh-lacZ*; magenta) and TTCs (*dSRF>GFP*; green) in control (Sucrose) or *Pe* infected midguts without (Damage) or with NAC (Damage + NAC), and following incubation in normal food during 32 hrs, post-infection (Recovery). Arrows indicate *trh-lacZ* signal within TTCs. Dotted boxes show magnified views of TTCs pointed by arrows. Scale bars: 50μm (main figure); 20μm (close up view). **f**, Quantification of *trh-lacZ* staining intensity relative to background within TTCs of midguts as in (e). <u>Statistics</u>: Student’s t test; n= number of TTCs from 12-21 posterior midguts per condition. **g**, Representative confocal images from control (Suc) or regenerating (*Pe*) adult posterior midguts from wild type animals or upon adult-specific *trh* overexpression (*trh*) or RNAi-driven knockdown (*trh^-IR^*) within TTCs (green). Scale bar: 100μm. **h**, **i**, Quantification of tracheal coverage (h) and PH3^+^ ISCs (i) in posterior midguts as in (g). <u>Statistics</u>: Two-way ANOVA, followed by Sidak’s multiple comparisons test; n= number of posterior midguts. **j**, Schematic representation of working model. Error bars: ±SEM; *p < 0.05, **p < 0.01, ***p < 0.001 ****p< 0.0001.

Here, we reveal a novel inter-organ communication program in *Drosophila*, involving the adult tracheal system and the midgut, which drives reciprocal adaptation of both tissues to sustain a robust regenerative response of the intestine to injury (Fig. 7j). Our results may reflect vasculature/stem cell interactions in the mammalian intestine and other self-renewing tissues.

## Discussion

The vasculature represents a prominent component of the gut microenvironment. However, its functional role in adult intestinal homeostasis remains largely unknown. Here, we report the cellular and molecular underpinnings of a novel inter-organ communication program between the adult *Drosophila* midgut and its closely associated vasculature-like tracheal tissue, which is fundamental to drive the regenerative response of ISCs following epithelial tissue damage.

ROS are key initiators of TTC remodelling and ISC proliferation, following intestinal damage by pathogenic infection (Fig. 7j). While our observations show a conserved phenomenology of TTCs/vasculature changes upon diverse intestinal insults and across species, it is highly conceivable that damage and species-specific molecular responses exist and that signals other than or in addition to ROS influence tracheal/vascular adaptations to intestinal damage.

HIF-1α/FGFR signaling drives tracheogenesis during development and in response to hypoxia following activation by FGF from target tissues^9, 34, 35^. As such, our findings suggest a repurposing of this developmental pathway during adult gut/tracheal crosstalk. However, the discovery of ROS-inducible angiocrine Bnl/FGF activating stem/progenitor cell FGFR signaling during adult intestinal regeneration is a novel finding from our study (Fig. 7j). FGF induces vascular endothelial cell differentiation in human intestinal organoids^53^ and acts as an angiocrine factor in various tumour settings^54^. This highlights the great degree of conservation between the *Drosophila* and mammalian system and raises the possibility of a conserved angiocrine role of FGF ligands in mammalian intestinal regeneration.

The DPP/BMP ligand has been reported as an angiocrine factor required to restrain stem cell proliferation in the adult *Drosophila* midgut^15^. However, knocking down *dpp* in adult TTCs did not affect basal stem cell proliferation as reported upon global ligand downregulation in the adult trachea^15^ (data not shown). Ligand compensation from main tracheal branches or the intestinal epithelium^55^ may explain discrepancies between our results and the published study.

In addition to the well-established role of oxygen, nutrition regulates TTC remodelling in the larval and adult *Drosophila* midgut^10^. In this context, a defined subset of enteric neurons influence TTC remodelling through the delivery of Insulin- and Vasoactive Intestinal Peptide-like neuropeptides^10^. Nutrient induced-tracheal remodelling involves activation of Insulin Receptor (InR) signaling within TTCs. We observed no requirement for InR or enteric neuronal activity in the plastic response of adult TTCs to intestinal damage (data not shown). Reciprocally, FGF/FGFR signaling does not appear to mediate nutrient dependent TTC remodelling^10^. These results suggest that molecular events driving tracheal tissue plasticity are diverse and highly dependent on the biological context and/or stimuli.

We identified two novel tracheal-intrinsic molecular mechanisms triggered in response to intestinal epithelial damage and necessary to induce TTC remodelling and ISC proliferation (Fig. 7j). One, involving upregulation of *Drosophila*IGF2 mRNA-binding protein (Imp) and its downstream target Myc (Fig. 7j). The other, requiring downregulation of the tracheal cell specification factor *trh* (Fig. 7j). Known functions of Imp had been restricted to the induction of neuronal remodelling and growth^48, 49^. Its mammalian orthologue, IGF2BP2, has been studied for its involvement in metabolic disease^56^ and its potential role in the vasculature remains to be addressed. Trh, homologous to mammalian NPAS3^57^, has been exclusively known for its requirement in the specification of tracheal cells from undifferentiated progenitors in the developing embryo^50–52^. Here, we present the first report for a role of Trh in terminally differentiated adult tracheal cells, which involves its unexpected downregulation. Emerging evidence suggests that adult tissues and cells, such as the intestine and neurons, lose differentiation markers and acquire ‘naïve’ or ‘foetal-like properties’ during the process of tissue regeneration^58–60^. Our work suggests the exciting possibility that this may also be the case for the adult vasculature.

The vasculature is a largely uncharacterized component of the adult intestinal niche. Vascularization of *in vitro* organ culture systems has been notoriously difficult, representing a major roadblock in the field of tissue engineering. As such, our *in vivo* findings may be of broad interest and impact to the vascular and intestinal research fields.

## Methods

### Fly stocks and rearing

A complete list of fly lines and full genotypes used in this study can be found in Supplementary Tables 5 and 6. In experiments using the *GAL4*/*GAL80^ts^* system, flies were crossed, F1 progenies reared, and adults aged for 5 days after eclosion at 18°C. Animals were then transferred to 29°C for 5-7 days to allow activation of most transgenes prior to phenotypic analysis. The exception was *bax*, which was overexpressed for only 3 days. If not carrying temperature sensitive transgenes, crosses and offspring were kept at 25°C. Overall, experimental animals were used 10-12 days following adult eclosion. Animals for experiments were maintained in food vials at low densities (10-15 flies per vial) and were transferred to fresh food every 2 days. Only adult posterior midguts from mated females were analysed in this study.

### Damage induced intestinal regeneration

Oral administration of *Pe*, Bleomycin or DSS was performed as previously described^21, 25, 61^ with minor modifications. Briefly, 10 day-old females of the desired genotypes were starved in empty vials for 2hrs followed by feeding with a 5% sucrose solution only (Sucrose), or Sucrose containing either *Pe* at OD 100, 25μg/ml Bleomycin (Sigma-Aldrich, Cat#B2434), or 3% DSS (Sigma-Aldrich, Cat#42867) applied on filter paper discs (Whatman). *Pe* infection was carried out for 16hrs, Bleomycin feeding was done for 1-day and DSS feeding lasted for 2-days, with fresh media applied each day.

### NAC treatment

Flies were placed in empty vials with a filter paper soaked with a 5% sucrose solution containing 20mM NAC (Sigma-Aldrich, Cat#A7250) for 24 hrs. Animals were then fed with either 5% sucrose + NAC or 5% sucrose + NAC + *Pe* (OD 100) for an additional period of 24 hrs.

### Mouse intestinal regeneration and IHC

*C57BL/6* mice were subject to whole body 10 Gy gamma-irradiation and the intestines were analysed 72 hrs post-irradiation, which represents the proliferative phase of the regenerative response to damage in the mouse small intestine^28^. Small intestines were isolated and flushed with tap water. 10× 1cm portions of small intestine were bound together with surgical tape and fixed in 4% neutral buffered formalin. Intestines from 3 mice per condition were used. 4 μm sections of formalin–fixed paraffin–embedded (FFPE) tissues were cut, mounted onto adhesive slides and incubated at 60°C overnight. Prior to staining, sections were dewaxed for 5 minutes in xylene followed by rehydration through decreasing concentrations of alcohol and final washing with H_2_O for 5 minutes. FFPE sections underwent heat–induced epitope retrieval in a Dako pre-treatment module. Sections were heated in Target Retrieval Solution High pH (Dako, K8004) for 20 minutes at 97°C before cooling to 65°C. Slides were removed and washed in Tris Buffered Saline with Tween (TbT) (Dako, K8000) and loaded onto a Dako autostainer link48 platform where they were stained with anti-CD31 antibody 1:75 (Abcam, ab28364) following standard IHC procedures. All animal work has been approved by a University of Glasgow internal ethics committee and performed in accordance with institutional guidelines under personal and project licenses granted by the UK Home Office to J.B.C (PPL PCD3046BA).

### Single terminal tracheal cell clones

To generate single cell clones of terminal tracheal cells, parental lines were allowed to mate for 2-3 days, after which adults were moved into new vials and F1 progenies where heat shocked in a water bath for 1 hr at 37°C. Adults of the correct genotype, emerging from the heat shocked animals, were selected and aged at 25°C for 10 days followed by feeding with Sucrose (control) or *Pe* for 16 hrs to cause intestinal damage and induce regeneration. Tissues were then dissected and processed for immunofluorescence staining and confocal imaging.

### Hypoxia treatment

Adult flies were aged at 25°C or 29°C in 21% O_2_ (Normoxia) at a density of 15-20 flies per vial. Then, animals were transferred overnight to 3% O_2_ (Hypoxia) in a Whitley Scientific H35 hypoxystation incubator.

### *Drosophila* immunohistochemistry

Immunohistochemistry was carried out as described previously^31^. The following antibodies were used: chicken anti-GFP 1:200 (Abcam, ab13970), mouse anti-PH3 1:100 (Cell Signaling, 9706), rabbit anti-Dcp1 1:100 (Cell Signaling, #9578S), rabbit anti-βgal 1:1000 (MP Biochemicals #559761), rabbit anti-DsRed 1:1000 (Clontech, #632496), mouse anti-Arm 1:3 (Hybridoma Bank, N2 7A1), mouse anti-Dlg 1:100 (Hybridoma Bank, 4F3), rabbit anti-p-Erk 1:100 (Cell Signaling, #9101), guinea pig anti-Myc 1:100 (gift from Gines Morata) and rabbit anti-Trh 1:100 (gift from M. Llimargas). Chitin Binding Protein (CBP 1:100; gift from M. Llimargas) was used to visualise all tracheal tissue. Alexa Fluor 488, 546 and 647 (Invitrogen) were used as secondary antibodies labels at 1:200 and 1:100 respectively. Guts were mounted in Vectashield anti-fade mounting medium for fluorescence with DAPI (Vector Laboratories, Inc) to visualize all nuclei.

### Image acquisition

#### Transmission Electron Microscopy

Guts were dissected under Schneider’s insect medium and were subsequently fixed in 2.5% glutaraldehyde in 0.1 M cacodylate buffer (pH 7.4) for 1hr at room temperature. Samples were rinsed repeatedly in 0.1M sodium cacodylate buffer, before fixation treatment in 1% Osmium Tetroxide/buffer for 1hr, followed by washing with dH20 for 30mins. Samples were then stained in 0.5% Uranyl Acetate/dH20 for 1hr (in the dark) and dehydrated by incubation in a graded series of Ethanol. Samples were subject to three subsequent incubations in Propylene Oxide followed by Epon 812 resin/Propylene Oxide (50:50) mix and left on a rotator overnight, followed by several incubations in pure Epon resin. Samples were then embedded into blocks and oven incubated at 60°C for 48hrs. Ultrathin sections (50-70nm thickness) were cut using a Leica Ultracut UCT. The sections produced were collected on Formvar coated 100 mesh copper grids and subsequently contrast stained in 2% Methanolic Uranyl Acetate for 5mins followed by Reynolds Lead Citrate for 5mins. Gut samples were viewed using a JEOL 1200 EX TEM and Images captured using a Cantega 2K x 2K camera and Olympus ITEM Software.

#### Confocal microscopy (Zeiss LSM 780)

Each image represents half-volume of the full posterior midgut (area comprised between hindgut and Copper Cell Region) and were acquired with 20x, 40x or 63x lenses using identical acquisition conditions for all samples from a given experiment. Images represent maximal intensity projection of a stable number of Z-Stacks and were processed with ImageJ and Carl Zeiss Zen 3.0 to adjust brightness and contrast.

#### Light microscopy (Axio observer Zeiss)

Adult guts were dissected in PBS, mounted in 100% glycerol and imaged immediately. Up to three pictures per posterior midgut were taken to cover most of the area. Images were taken at their most apical plane to best detect TTCs with a 20x lens. Images from mouse intestinal samples (Fig. 1n) were acquired under a 20x lens using this microscope and were processed with ImageJ.

### Quantifications in the adult posterior midgut

#### Quantification of ISC proliferation

Antibodies against Phosphorylated Histone H3 (PH3) were used to detect ISC proliferation in the adult midgut. The total number of PH3^+ve^ cells per posterior midgut was quantified manually upon visual inspection using an Olympus BX51 microscope. The number of midguts analysed (n) for each experiment are indicated in the figures.

#### Quantification of tracheal coverage from immunofluorescence images

Unless otherwise noted, confocal images of *dSRF>GFP* expressing midguts were used and tracheal values provided were obtained from quantification of tile scan images of the entire R4-R5 posterior midgut regions, acquired with a 20x lens. Values of tracheal coverage represent pixel per area and were obtained from maximum intensity Z-projections. Pictures were individually processed on ImageJ as follows: 1) maximum intensity projection from Z stacks were produced; 2) area of interest was cropped to eliminate Malpighian tubules, hindgut and copper cell region; 3) “threshold” was adjusted to ensure the detection of most of terminal tracheal branches; 4) function “skeletonize” was applied to generate a skeleton of the tracheal network; 5) maximum intensity of this skeleton was measured (Extended data Fig. 1a). The number of posterior midguts analysed (n) for each experiment are indicated in the figures.

#### Quantification of tracheal branching form light-microscopy images

Acquired images were blindly scored using a 1 to 5 scoring system (Extended Data Fig. 3a, c). The custom ImageJ macro used for blind tracheal scoring is available at “<u>Blind_scoring.ijm</u>”. Number of midguts analysed (n) for each experiment are indicated in the figures.

#### Quantification of total branches per TTC and TTC ramifications

Maximum intensity projections from confocal Z stacks were used. The number of primary, secondary and tertiary branches derived from individual TTCs was assessed (Fig. 1i). Due to the intrinsic complexity of the tracheal, it is difficult to unambiguously assign cellular extensions/branches to a single TTC. To circumvent this issue, we counted tracheal branches starting from a TTC body and defined the end of a TTC extension when it touched the body of another TTC. Additionally, the generation of single TTC clones allowed us to unambiguously quantify the total number of branches from individual TTC, which we did manually (Fig. 1f, h). These two approaches led to same outcome. Number of TTC (n) and midguts analysed for each condition are indicated in the corresponding figures and figure legends.

#### Quantification of total tracheal length

We used the plugin “NeuronJ” from ImageJ to quantify the total length of all branches emerging from a TTC (Fig. 1k). Number of TTC (n) and midguts analysed for each condition are indicated in the corresponding figures and figure legends.

#### Quantification of TTC nuclei

Confocal images of posterior midguts from animals expressing *dSRF^ts^>RedStinger* to label TTC nuclei were used. The custom ImageJ macro used for quantifying the number of TTC nuclei is available at <u>“dSrf_pH3_overlap_for Jessica.ijm”</u>. This macro was also used to quantify the number of TTC nuclei positive for PH3^+^ staining (Extended data Fig. 1g). Number of midguts analysed (n) for each condition is indicated in the corresponding figure and figure legend.

#### Quantification of posterior midgut area

Midgut tissue was visualized by DAPI staining and the posterior midgut area (length x width) was measured with ImageJ (Extended Data Fig. 1c, d). Number of midguts analysed (n) for each condition is indicated in the corresponding figure and figure legend.

#### Quantification of lacZ reporters

Antibodies against β-galactosidase were used to detect *lactate dehydrogenase*-, *Delta*-, *breathless*-, *bs/dSRF*-, *branchless*- and *trachealess-lacZ* reporters. Pictures were taken with confocal microscopy and staining was quantified using ImageJ. For each gut quantified, the background staining signal was subtracted from the total signal of β-galactosidase detected in TTCs, ISCs/EBs or ECs. This value was then divided by the background signal to normalize the data. Number of cells (n) and midguts analysed for each condition are indicated in the corresponding figures and figure legends.

#### Quantification of cell death

Antibodies against Dcp1 were used to assess cell death in posterior midguts. Pictures were taken with confocal microscopy and Dcp1 staining intensity was measured relative to the surface of the gut area analysed. Number of midguts analysed (n) for each experiment are indicated corresponding in the figure and figure legend.

#### Quantification of pERK, Imp::GFP, Myc and Trachealess staining

Midguts stained with antibodies to detect these proteins included a methanol fixation step between the PFA fixation and PBST washing steps of the standard protocol, as described previously^40^. Images were acquired with confocal microscopy and staining was quantified using ImageJ. For each gut quantified, the background staining signal was subtracted from the total antibody signal within DAPI positive cells. This value was then divided by the background signal in order to normalize the data. The number of cells (n) and midguts analysed for each condition are indicated in the corresponding figures and figure legends.

### Statistical analysis

Unless otherwise noted, in all figures: NS, not significant (p>0.05); *p < 0.05, **p < 0.01, ***p < 0.001 ****p< 0.0001. Prism 6 software (GraphPad) was used for statistical analyses. A two-tailed Student’s t-test was conducted to compare two sample groups. To test for significance in larger groups, two-way ANOVAs, corrected for multiple comparisons using Sidak’s statistical hypothesis testing, respectively, were used. P values less than 0.05 were considered significant. Information on sample size and statistical tests used for each experiment are indicated in figure legends. Unless otherwise noted, experiments represent a minimum of 2 independent biological replicates. Please see reporting summary for further details.

### qRT-PCR

Trizol (Invitrogen) was used to extract total RNA from 30 midguts per biological replicate. cDNA synthesis was performed using the High-Capacity cDNA Reverse Transcription Kit (Applied Biosystems). QuantiNova SYBR Green (Qiagen) was used for qPCR. Data were extracted and analysed using QuantStudio and Prism 6.0. Data from 5 biological replicates is presented as the mean fold change with standard error. Expression of target genes was measured and normalized to *gapdh1* or *act5c* using standard curves. Primer sequences can be found in Supplementary Table 4.

### Targeted DamID (TaDa), library preparation, sequencing and data analysis

*dSRF-GAL4; tub-gal80^ts^ (dSRF^ts^>)* animals were crossed to *UAS-LT3-Dam* or *UAS-LT3-Dam-Pol II* animals at 18°C. F1 progeny were collected every 48 hrs and aged for a further 7 days at 18°C before transferring to 29°C to induce adult restricted Dam protein expression for 24 hrs. During the last 16 hrs at 29°C, flies were fed a Sucrose or Sucrose + *Pe* solution. 60 midguts per condition per biological replicate were dissected in cold PBS and stored at −80°C. Methylated DNA fragments were isolated and next generation sequencing libraries were prepared as described previously^45^. Sequencing data from TaDa experiments were processed using a previously described pipeline^62^ and mapped to release 6.03 of the *Drosophila* genome. Transcribed genes were annotated for Pol II binding data using a custom Perl script^46^ (available at https://github.com/tonysouthall/Dam-RNA_POLII_analysis) and release 6.11 of the annotated *Drosophila* genome. Genes with significant RNA pol II binding were identified based on meeting a threshold of 1% FDR and > 0.2 log2 ratios. Briefly, a log2 ratio of the Dam-RNA Pol II read counts over control Dam-only read counts is calculated after quantile normalisation^62^ and if this ratio is higher than 0.2, then we would determine that this gene has significant RNA Pol II binding (Supplementary Table 1). Significance was assigned based on the signal from multiple GATC fragments and using a very stringent pipeline as a transcript had to have a false discovery ratio (FDR) of less than 1% in both replicates to be called significant.

Gene Ontology (GO) term analysis was performed using the R package ClusterProfiler^63^ to search for enriched GO terms (Supplementary Table 2). Three independent biological replicates were originally processed for each condition (Sucrose and *Pe*). However, after visual inspection of the sequencing tracks, one replicate from each condition was excluded from the analysis, due to poor DNA sample quality and unreliable sequencing data. Scattered plots showing the correlation between samples for each condition are provided (Extended data Fig. 7a).

## Data availability

TaDa sequencing data is available through the University of Glasgow institutional repositories *DOI: 10.5525/gla.researchdata.994 (2020)*. The entire set of raw data supporting this study will be made accessible through the same repository prior to publication.

## Acknowledgements

J.P. and J.B.C. are funded by a Wellcome Trust and Royal Society Sir Henry Dale Fellowship (Grant Number 104103/Z/14/Z; J.B.C.) and a Wellcome Trust Institutional Strategic Support Fund (ISSF) — Excellence and Innovation Catalyst Award to J.B.C. J.P. was partly funded by a BBSRC - Flexible Talent Mobility Account (FTMA) Award (BB/R506576/1). Y.Y. is supported by CRUK core funding to the CRUK Beatson Institute (A17196). T.D.S and G.N.A. were funded by a Wellcome Trust Investigator grant (104567; T.D.S.) and a BBSRC grant (BB/P017924/1; T.D.S. and G.N.A.)

We are thankful to Marta Llimargas, Markus Affolter, Irene Miguel-Aliaga, Andrea Brand, Cedric Polesello, Alessandro Scopelliti, Pablo Wappner, Ilan Davis, Florence Besse, Hugo Stoker, Gines Morata and Fisun Hamaratoglu Dion for generously sharing reagents and fly lines. We thank the Kyoto, Vienna and Bloomington Drosophila Stock Centres and the Drosophila Studies Hybridoma Bank for fly stocks and antibodies. We thank Colin Nixon (Beatson CRUK histology service) for IHC of mouse intestinal samples, Elaine McKenzie for help and training on the use of the hypoxia chamber, Margaret Mullin for assistance with TEM and David Strachan, John Halpin and Robert Insall (Beatson CRUK) for support with image quantification. We thank David McGuinness and Julie Galbraith (Glasgow Polyomics) for sequencing samples for TaDa and Rhoda Stefanatos for advice with RT-qPCRs. We thank Maté Naszai for help with the creation of custom ImageJ macros and Lynsey Carroll for providing mouse intestinal samples. We thank Jean-Philippe Parvy and multiple members of the Cordero lab for scientific discussion and advice on the project.

## Author contributions

J.P designed and carried out most experiments and analysed and interpreted the data. Y.Y. provided technical support throughout the study and perform RT-qPCRs. J.P. G.A, T.S and J.B.C. analysed the TaDa data. J.B.C conceived the project, designed experiments, analysed the data and supervised the study. J.P and J.B.C wrote the paper with contributions from the rest of the authors.

---

**Extended data Fig. 1.**
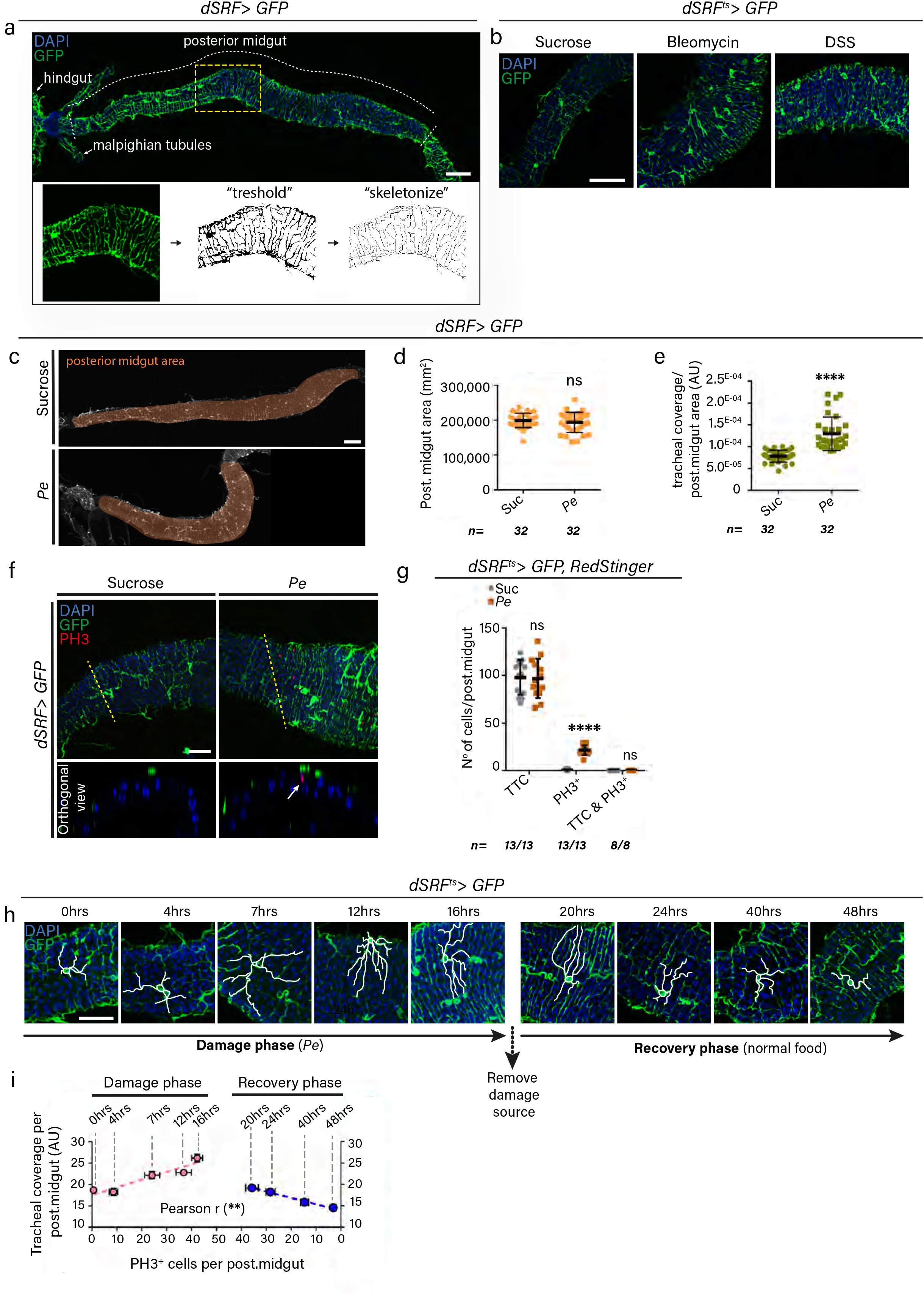
Characterization and quantification of adult tracheal remodelling following intestinal damage. **a, (top)**, Representative confocal image of adult posterior midgut and associated TTCs (green). **(bottom)**, illustration of the different steps followed for the quantification of gut tracheal coverage as explained in Methods. Scale bar: 100μm. Box in top panel highlights the area shown in the bottom panels **b,** Representative confocal images of TTCs (green) from posterior midguts of control animals (Suc) or animals fed with Bleomycin or DSS. Scale bar: 50μm. **c**, Representative confocal images of control (Suc) or damaged (*Pe*) adult posterior midguts (shaded in brown). Scale bar: 100μm. **d**, **e**, Quantification of posterior midgut area (d) and ratio of tracheal coverage over adult posterior midgut area (e). **f**, Representative confocal images of TTCs (green) from control (Suc) or *Pe* damaged adult posterior midguts stained with anti-PH3 to label proliferating ISCs (red). Bottom panels represent orthogonal views of the midguts shown in top panels. Scale bar: 50μm. **g,** Quantification of individual or combined TTC nuclei and PH3^+^ cells in control or *Pe* infected midguts. **c-g**, <u>Statistics</u>: Student’s t test; n= number of posterior midguts; error bars ±SEM. ****p< 0.0001. **h**, Confocal images of adult TTCs assessed at the indicated time points during and after intestinal damage. White lines trace individual TTCs. Scale bar: 50μm. **i**, Correlation graph between TTC coverage and ISC proliferation for each of the time points and conditions presented in (h). <u>Statistics</u>: Pearson’s correlation coefficient (r)=0.9660 and 0.9962; **p=0.0075 and 0.0038 in damage and recovery phases, respectively.

**Extended data Fig. 2.**
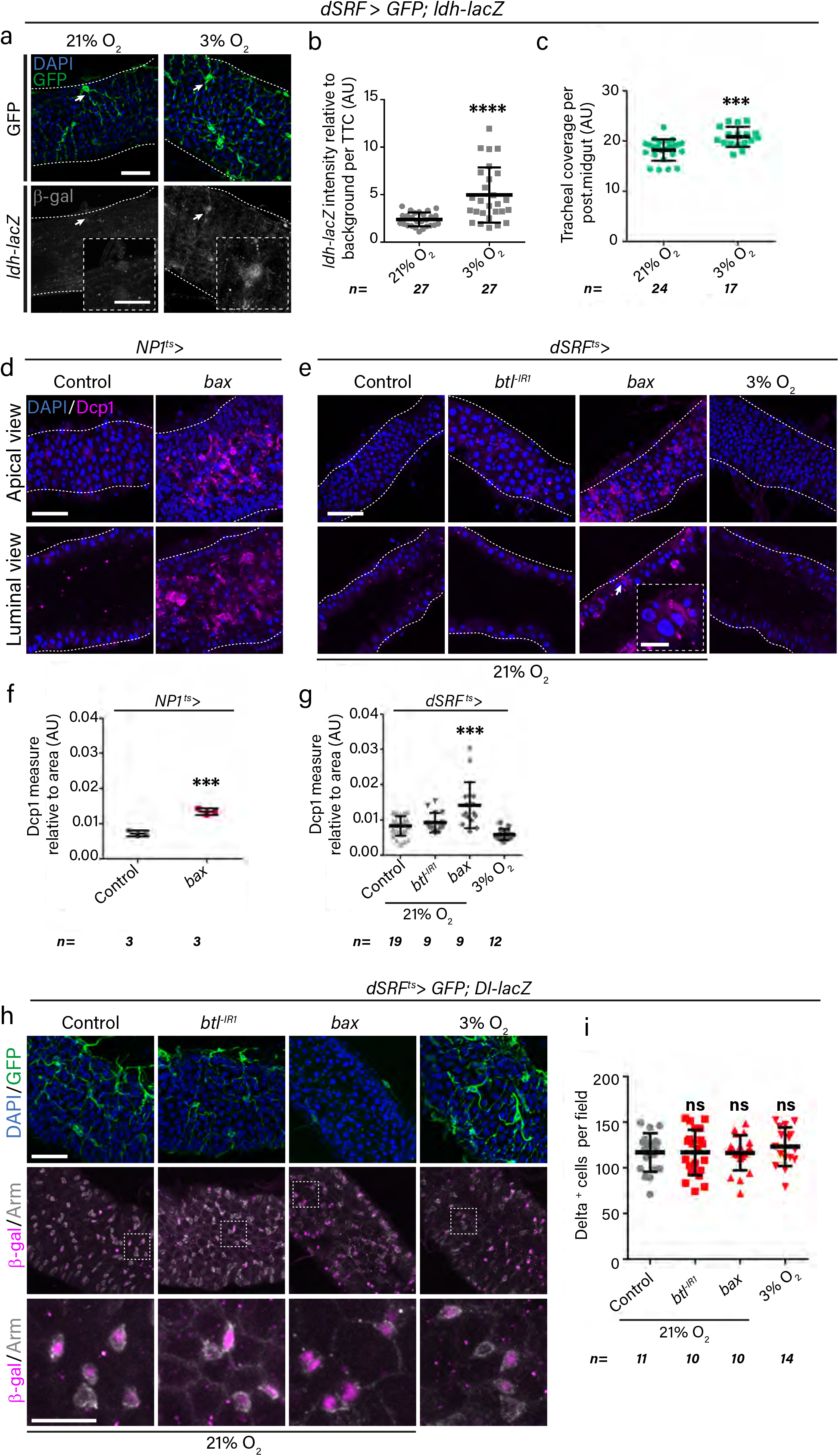
Hypoxia or impaired TTC remodelling does not affect ISC survival. **a,** Confocal images of TTCs (green) and the HIF-1α/Sima activity reporter *ldh-lacZ* (grey) in adult posterior midguts from animals in normoxia (21% O_2_) or subjected to 24 hrs of hypoxia (3% O_2_). Dotted boxes show a magnified view of TTCs pointed by arrows. Scale bars: 50μm (main figure); 20μm (close up view). **b**, Quantification of *ldh-lacZ* staining intensity within TTCs, relative to background, in midguts as in (a). <u>Statistics</u>: Student’s t test; n= number of TTC from 9 posterior midguts per condition. **c**, Quantification of tracheal coverage in adult posterior midguts as in (a). <u>Statistics</u>: Student’s t test; n= number of posterior midguts. **d, e**, Confocal images of adult posterior midguts stained with anti-Dcp1 (magenta) to visualize cell death in control animals; upon adult-specific *bax* overexpression in ECs (d); animals subjected to hypoxia or to the indicate genetic TTC disruptions (e). **d, e**, Upper panels (apical tissue views); lower panels (longitudinal sections showing intestinal tube lumen). Scale bar: 50μm. The high level of Dcp1 staining in the lumen of midguts subject to *bax* overexpression in ECs (d) corresponds to many delaminating/dying cells. Dotted box in (e) shows a magnified view of an apoptotic EC pointed by arrow in main figure and identified by its large nuclei and Dcp1 staining. Scale bars: 50μm (main figure); 20μm (close up view). **f**, **g,** Quantification of Dcp1 staining in midguts as in (d, e). <u>Statistics</u>: Student’s t test; n= number of posterior midguts; **h,** Confocal images of TTCs (green) and ISCs detected with a *Delta-lacZ* reporter (magenta) and anti-Armadillo (Arm) staining (grey) in midguts as in (e). Dotted boxes in middle panels indicate the magnified areas in the lower panels. Scale bars: 50μm (main figure); 20μm (close up view). **i**, Quantification of the number of *Delta-lacZ* positive cells in midguts as in (h). <u>Statistics</u>: Student’s t test; n= number of posterior midguts. Error bars: ±SEM; *p < 0.05, **p < 0.01, ***p < 0.001 ****p< 0.0001.

**Extended data Fig. 3.**
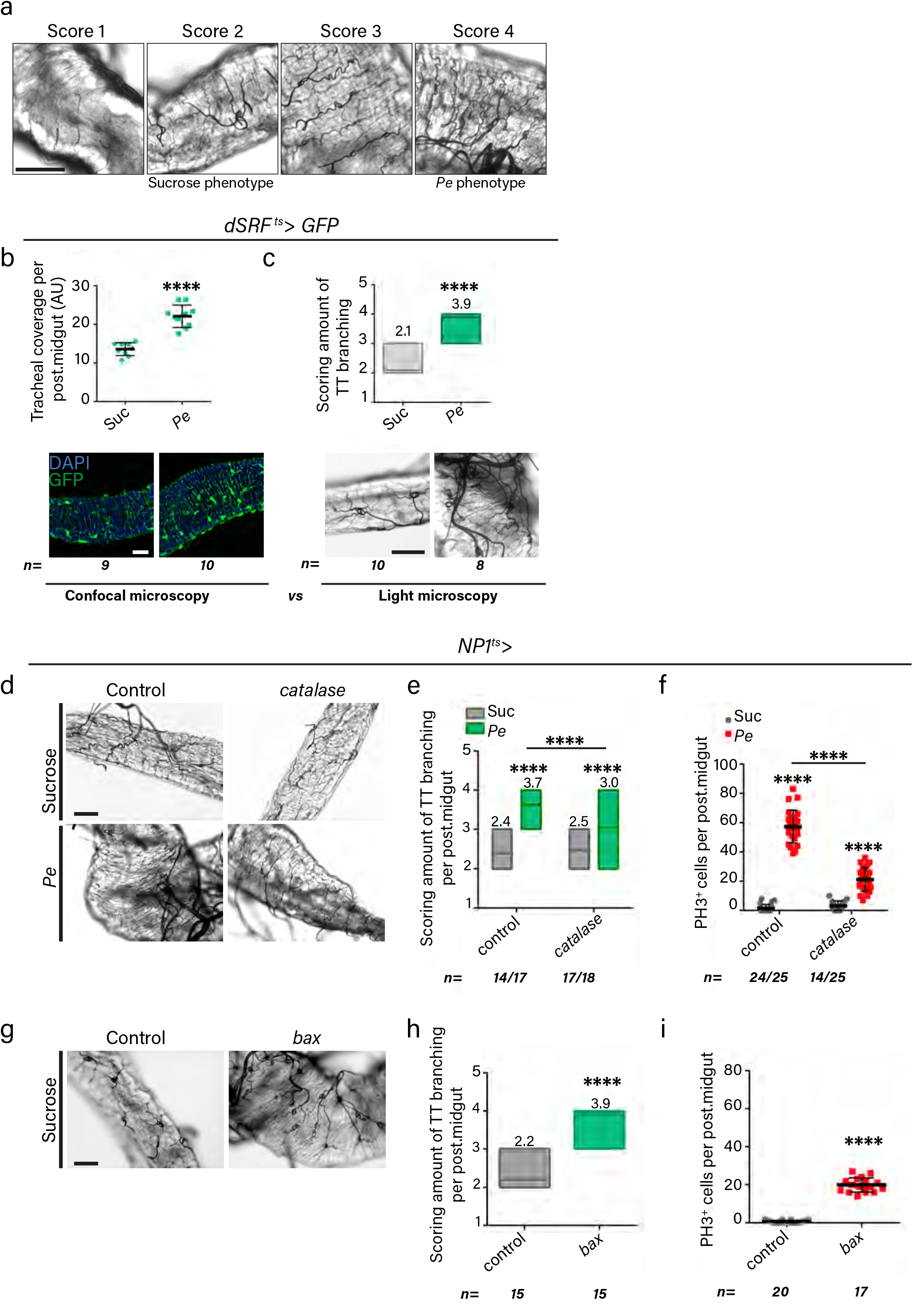
Gut associated TTCs are responsive to local signals from the damaged intestinal epithelium. **a,** Examples of tracheal branching levels assigned to each of the scores used for the quantification of tracheal coverage from light microscopy images. Scale bar: 50μm. **b**, **c (top panels)**, Quantification of tracheal branching from confocal (b) and brightfield images (c) of control (Suc) or *Pe* damaged posterior midguts. **b**, **c (bottom panels)** Representative confocal (b) or brightfield images (c) from midguts as in top panels. Scale bars: 50μm. **d**, Representative brightfield images from control (Suc) or *Pe* damaged posterior midguts from wild type animals or upon adult-specific *catalase* overexpression within ECs. Scale bar: 50μm. **e**, **f**, Scoring of tracheal branching (e) and quantification of PH3^+^ ISCs (f) in posterior midguts as in (d). <u>Statistics</u>: Two-way ANOVA followed by Sidak’s multiple comparisons test; n= number of posterior midguts quantified. **g**, Representative brightfield images of posterior midgut from control animals or upon adult-specific overexpression of *bax* in ECs. Scale bar: 50μm. **h**, **i**, Quantification of tracheal branching (h) and PH3^+^ ISCs (i) in posterior midguts as in (g). **b, c and g-i,** <u>Statistics</u>: Student’s t test; n= number of posterior midguts. Error bars: ±SEM; *p < 0.05, **p < 0.01, ***p < 0.001 ****p< 0.0001.

**Extended data Fig. 4.**
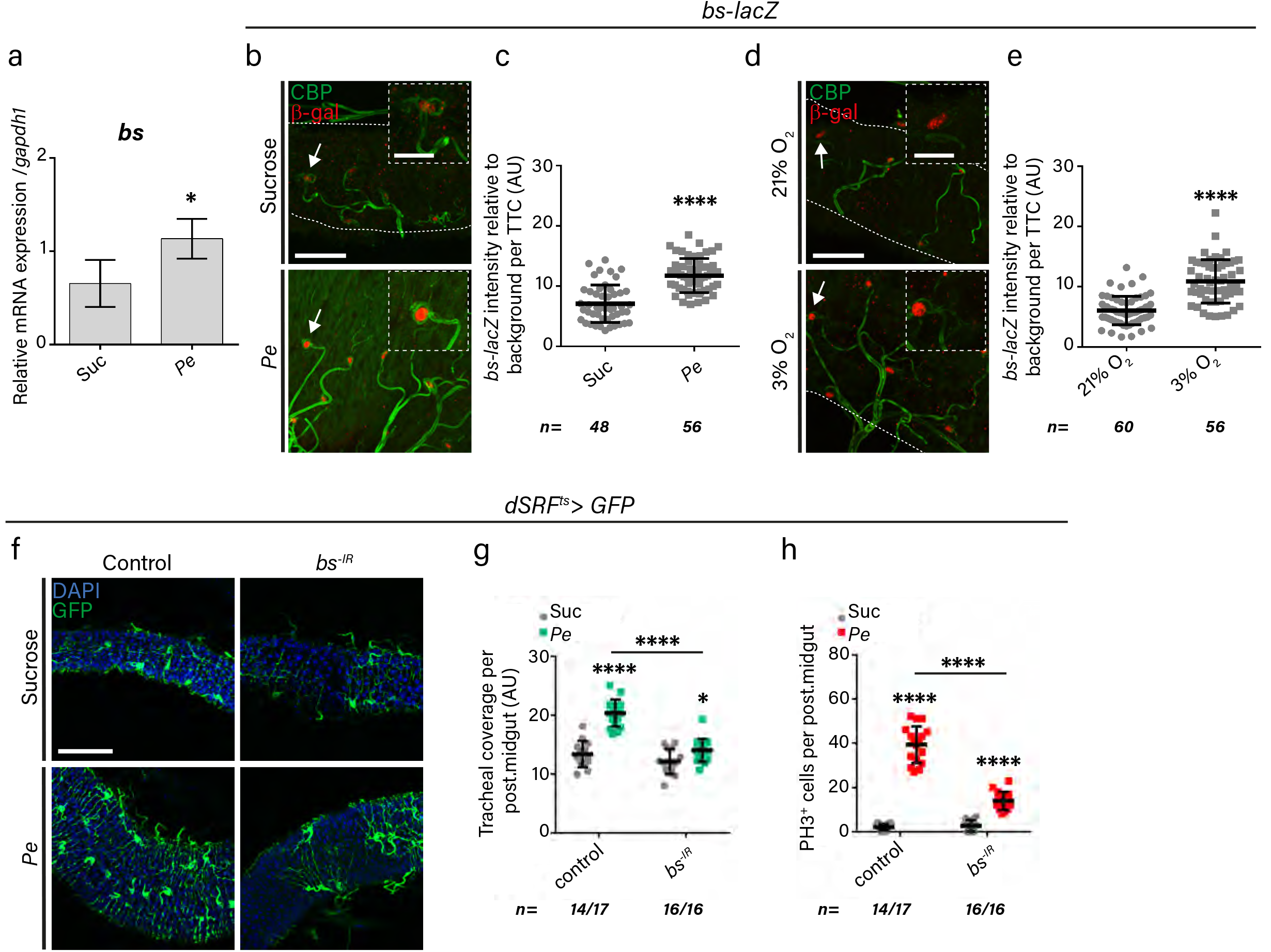
*blistered*/*dSRF* expression is upregulated in adult TTCs following intestinal damage or hypoxia and it regulates damage induced TTC remodelling and ISC proliferation. **a**, qRT-PCR for *blistered (bs)/dSRF* mRNA expression from adult midguts and associated tracheal tissue, in control (Sucrose) or *Pe* treated midguts. <u>Statistics</u>: Data are represented as mean ± SEM of five biological replicates; Student’s t test. **b**, Representative confocal images of *bs-lacZ* reporter expression (red) in control (Sucrose) and regenerating (*Pe*) adult posterior midguts stained with Chitin Binding Protein (CBP, green) to visualise all tracheal tissue. Dotted boxes show a magnified view of TTCs pointed by arrows. Scale bars: 50μm (main figure); 20μm (close up view). **c**, Quantification of *bs-lacZ* staining intensity within TTCs in posterior midguts as in (b). **d**, Representative confocal images of *bs-lacZ* reporter expression (red) in adult posterior midguts from animals housed in normoxia (21% O_2_) or subjected to 16 hrs of hypoxia (3% O_2_) and stained with CBP (green). Dotted boxes show a magnified view of TTCs pointed by arrows. Scale bars: 50μm (main figure); 20μm (close up view). **e**, Quantification of *bs-lacZ* staining intensity within TTCs in posterior midguts as in (d). **b-e,** <u>Statistics</u>: Student’s t test; n= number of TTC from 12-15 posterior midguts per condition. **f**, Confocal images of control (Sucrose) and damaged (*Pe*) adult posterior midguts from wild type animals or following *RNAi*-driven adult-specific *bs* knockdown (*bs^-IR^*) within TTCs. Scale bar: 100μm. **g**, **h**, Quantification of tracheal coverage (g) and PH3^+^ ISCs in posterior midguts as in (f). <u>Statistics</u>: Two-way ANOVA followed by Sidak’s multiple comparisons test; n= number of posterior midguts quantified. Error bars: ±SEM; *p < 0.05, **p < 0.01, ***p < 0.001 ****p< 0.0001.

**Extended data Fig. 5.**
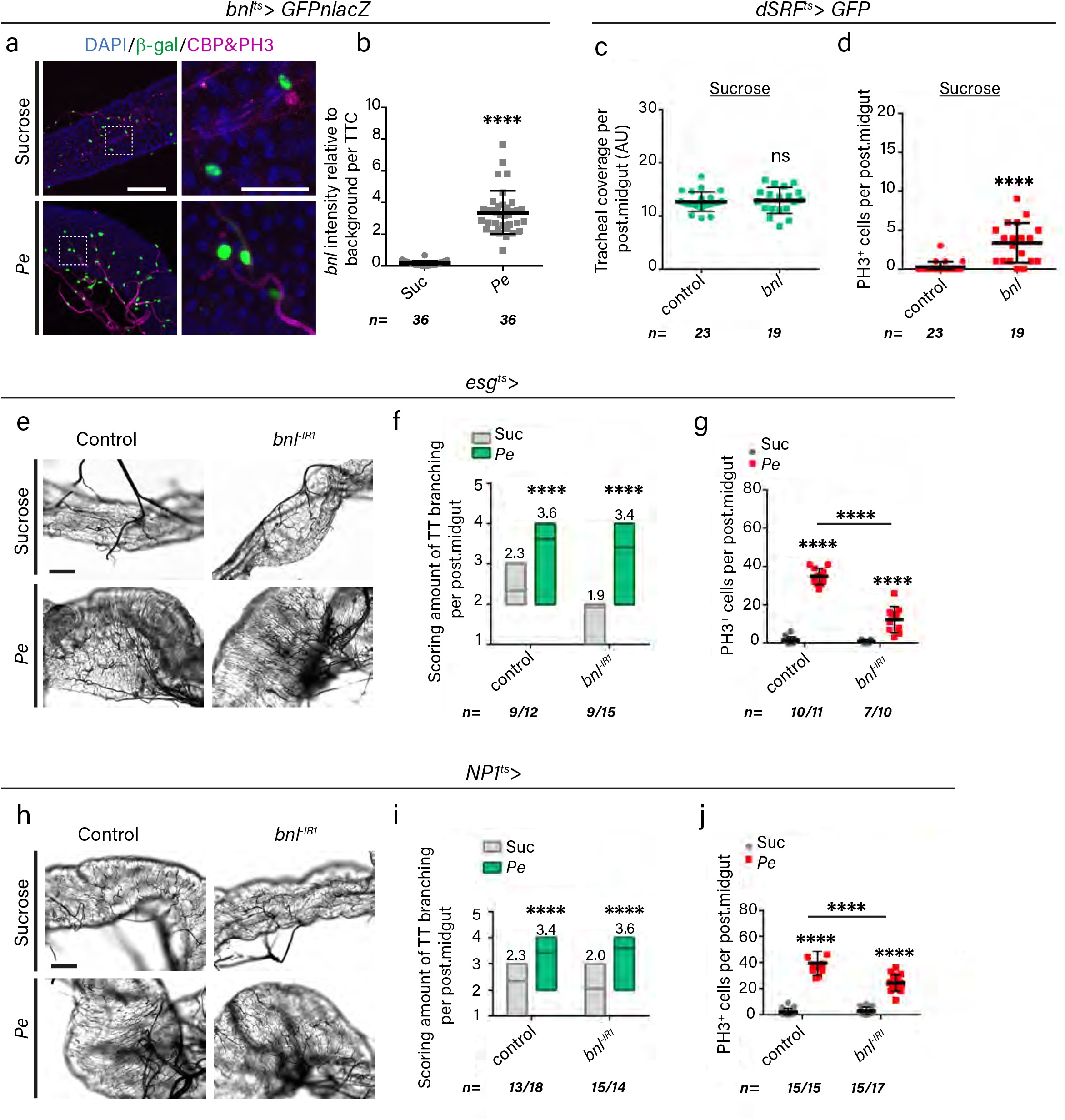
Multiple sources of Bnl individually contribute to regenerative ISC proliferation in the adult *Drosophila* midgut, independently of tracheal remodelling. **a**, Representative confocal images of *FGF/bnl* reporter expression *bnl>GFPnlacZ* (green) in control (Sucrose) and regenerating (*Pe*) adult posterior midguts stained with CBP (magenta) to visualise all tracheal tissue and PH3 (magenta). Dotted boxes in left panels indicate the area magnified in the right panels. Scale bars: 100μm (main figure); 40μm (close up view). **b**, Quantification of *bnl>GFPnlacZ* staining intensity relative to background within TTCs in posterior midguts as in (a). <u>Statistics</u>: Student’s t test; n= number of TTC from 12 posterior midguts per condition. **c, d** Quantification of tracheal coverage (c) and PH3^+^ ISCs (d) from wild type animals or upon adult-specific *bnl* overexpression within TTCs. <u>Statistics</u>: Student’s t test; n= number of posterior midguts. **e**, Representative brightfield images from control (Suc) or *Pe* damaged posterior midguts from wild type animals or following RNAi-driven adult-specific *bnl* knockdown (*bnl^-IR^*) within ISCs/EBs. Scale bar: 50μm. **f**, **g**, Quantification of tracheal branching (f) and PH3^+^ ISCs (g) in posterior midguts as in (e). **h**, Representative brightfield images from control (Suc) or *Pe* damaged posterior midguts from wild type animals or upon adult-specific *bnl* knockdown (*bnl^-IR^*) within ECs. Scale bar: 50μm. **i**, **j**, Quantification of tracheal branching (i) and PH3^+^ ISCs (j) in posterior midguts as in (h). **e-j,** <u>Statistics</u>: Two-way ANOVA followed by Sidak’s multiple comparisons test; n= number of posterior midguts quantified. Error bars: ± SEM; *p < 0.05, **p < 0.01, ***p < 0.001 ****p< 0.0001.

**Extended data Fig. 6.**
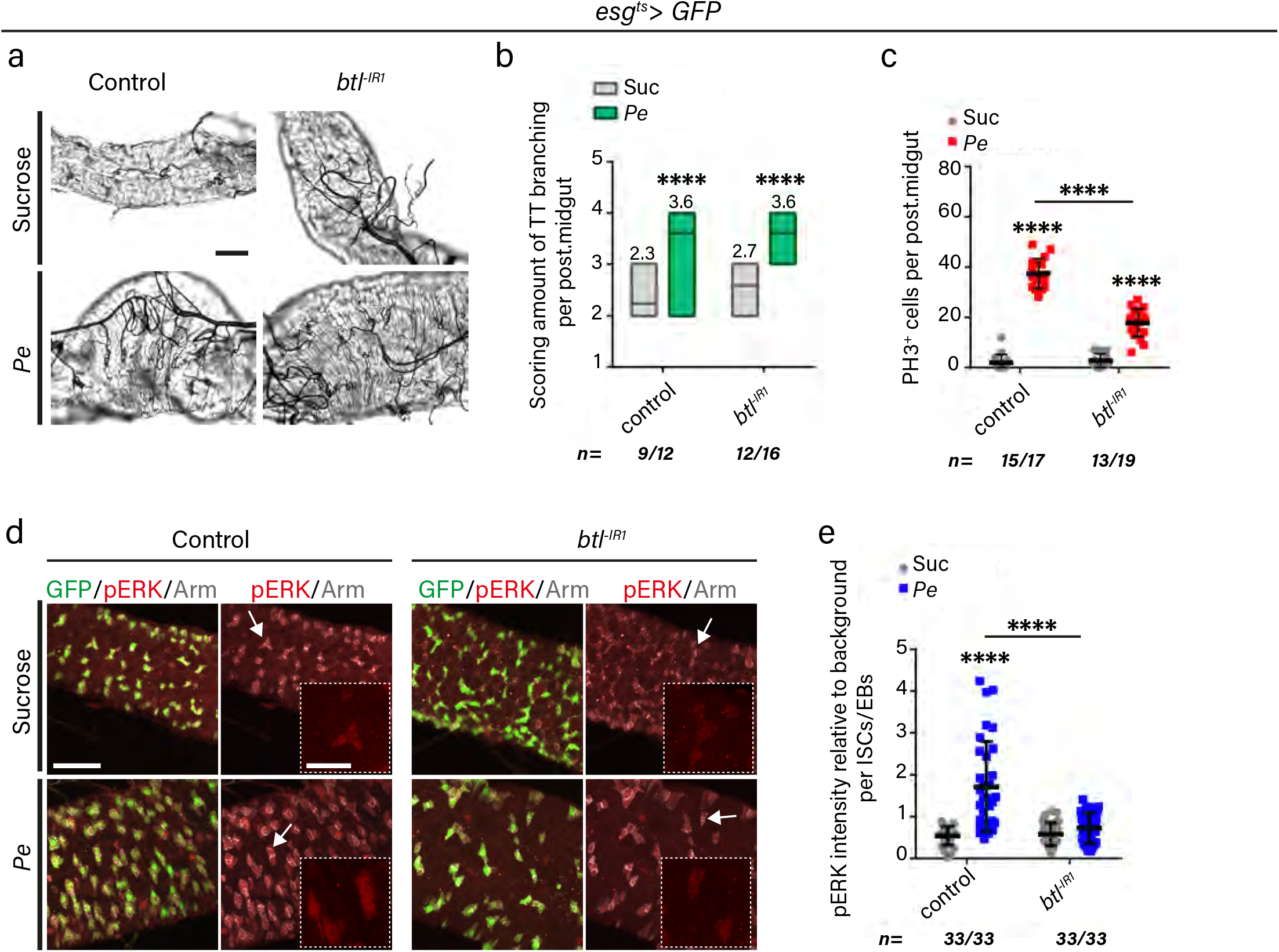
Individual sources of gut-derived Bnl are redundant for TTC remodelling but required for damage induced ISC proliferation. **a,** Representative brightfield images of control (Sucrose) or regenerating (*Pe*) adult posterior midguts from wild type animals or upon RNAi-driven adult-specific *btl* knockdown (*btl^-IR^*) within ISCs/EBs. Scale bar: 50μm. **b**, **c**, Quantification of tracheal branching (b) and PH3^+^ ISCs (c) in posterior midguts as in (a). <u>Statistics</u>: Two-way ANOVA followed by Sidak’s multiple comparisons test; n= number of posterior midguts. **d**, Representative confocal images of activated MAPK (pERK) staining (red), Arm (grey) and ISCs/EBs (*esg>GFP*, green) in control (Sucrose) or regenerating (*Pe*) adult posterior midguts from wild type animals or upon RNAi-driven adult-specific *btl* knockdown (*btl^-IR^*) within ISCs/EBs. Dotted boxes show a magnified view of the ISCs/EBs pointed by arrows. Scale bars: 50μm (main figure); 40μm (close up view). **e**, Quantification of pERK staining intensity relative to background within ISCs/EBs in posterior midguts as in (d). <u>Statistics</u>: Two-way ANOVA followed by Sidak’s multiple comparisons test; n= number of ISCs/EBs from 11 posterior midguts per condition. Error bars: ±SEM; *p < 0.05, **p < 0.01, ***p < 0.001 ****p< 0.0001.

**Extended data Fig. 7.**
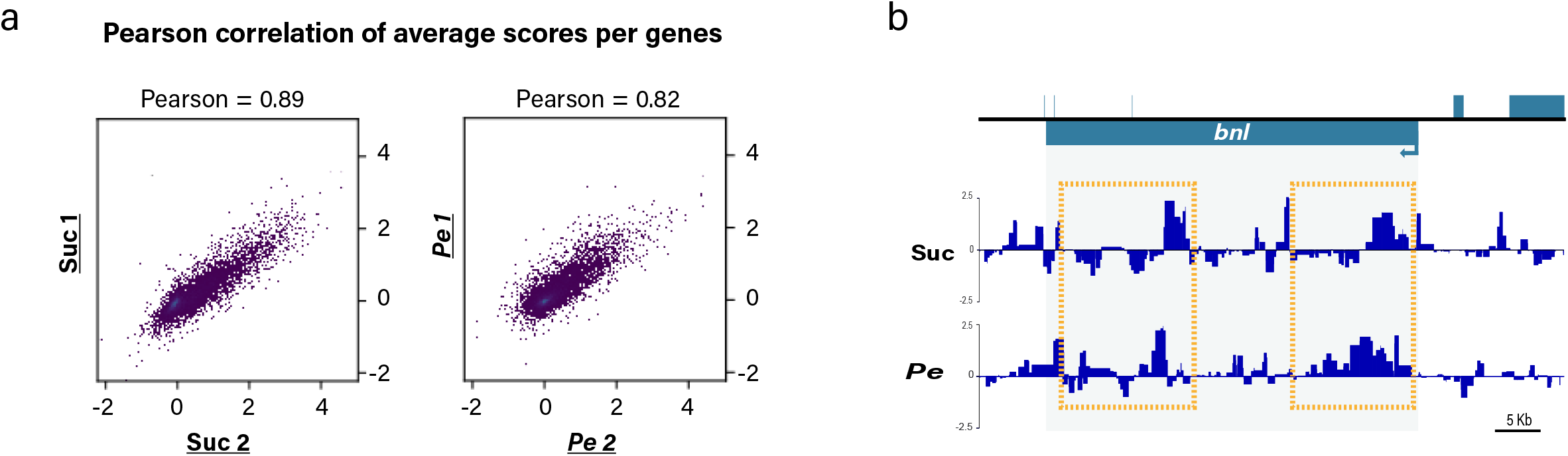
Genome wide RNA pol II binding profile within adult TTC by targeted DamID (TaDa). **a,** Scatterplots indicating correlation between TaDa replicates for each of the conditions used in this study. Significant correlation is observed between replicates of each condition. Each data point represents the average score for each gene (log2 Dam-pol II/ Dam-only). **b**, RNA Pol II binding profile to *bnl* in TTCs from control (Suc) and damaged (*Pe*) adult midguts. Boxes highlight areas with most prominent differences in RNA Pol II binding.

